# Heterochromatin diversity modulates genome compartmentalization and loop extrusion barriers

**DOI:** 10.1101/2021.08.05.455340

**Authors:** George Spracklin, Nezar Abdennur, Maxim Imakaev, Neil Chowdhury, Sriharsa Pradhan, Leonid Mirny, Job Dekker

## Abstract

Two dominant processes organizing chromosomes are loop extrusion and the compartmental segregation of active and inactive chromatin. The molecular players involved in loop extrusion during interphase, cohesin and CTCF, have been extensively studied and experimentally validated. However, neither the molecular determinants nor the functional roles of compartmentalization are well understood. Here, we distinguish three inactive chromatin states using contact frequency profiling, comprising two types of heterochromatin and a previously uncharacterized inactive state exhibiting a neutral interaction preference. We find that heterochromatin marked by long continuous stretches of H3K9me3, HP1α and HP1β correlates with a conserved signature of strong compartmentalization and is abundant in HCT116 colon cancer cells. We demonstrate that disruption of DNA methyltransferase activity dramatically remodels genome compartmentalization as a consequence of the loss of H3K9me3 and HP1 binding. Interestingly, H3K9me3-HP1α/β is replaced by the neutral inactive state and retains late replication timing. Furthermore, we show that H3K9me3-HP1α/β heterochromatin is permissive to loop extrusion by cohesin but refractory to CTCF, explaining a paucity of visible loop extrusion-associated patterns in Hi-C. Accordingly, CTCF loop extrusion barriers are reactivated upon loss of H3K9me3-HP1α/β, not as a result of canonical demethylation of the CTCF binding motif but due to an intrinsic resistance of H3K9me3-HP1α/β heterochromatin to CTCF binding. Together, our work reveals a dynamic structural and organizational diversity of the inactive portion of the genome and establishes new connections between the regulation of chromatin state and chromosome organization, including an interplay between DNA methylation, compartmentalization and loop extrusion.

**Highlights:**

- Three inactive chromatin states are distinguishable by long-range contact frequencies in HCT116, respectively associated with H3K9me3, H3K27me3 and a H3K9me2 state with neutral contact preferences.
- H3K9me3-HP1α/β heterochromatin has a high degree of homotypic affinity and is permissive to loop extrusion but depleted in extrusion barriers.
- Disrupting DNA methylation causes widespread loss of H3K9me3-HP1α/β and dramatic remodeling of genome compartmentalization.
- H3K9me3-HP1α/β is replaced by the neutral inactive state, which gains CTCF loop extrusion barriers and associated contact frequency patterns.
- DNA methylation suppresses CTCF binding via two distinct mechanisms.

## Introduction

Chromosomes are non-randomly organized inside the nucleus (Bolzer et al., 2005). Chromosome organization is associated with vital cellular processes including transcriptional regulation, epigenetic modification, DNA repair, and replication timing (McCord et al., 2020). High-throughput chromosome conformation capture (Hi-C) is a next-generation sequencing technique that can identify many features of chromosome architecture via proximity ligation. Patterns in Hi-C contact frequency maps reflect paradigms of chromosome organization such as chromosome territoriality, compartmentalization, and loop extrusion (reviewed in (Mirny et al., 2019; Oudelaar & Higgs, 2021)).

The best characterized chromosome organizing process is loop extrusion, which leads to the formation of topologically associating domains (TADs) and associated features (stripes and dots) in Hi-C maps from mammalian cells (Fudenberg et al., 2016; Sanborn et al., 2015). During interphase, cohesin complexes act as motors to extrude progressively growing chromatin loops. In vertebrates, the insulator protein CTCF serves as a directional barrier that halts loop extruding cohesin (de Wit et al., 2015; Fudenberg et al., 2016; Nora et al., 2020; Rao et al., 2014; Sanborn et al., 2015; Vietri Rudan et al., 2015). Independent of loop extrusion, chromosomes are also spatially compartmentalized: active chromatin is generally spatially segregated from inactive chromatin with euchromatin located more centrally and heterochromatin more peripherally in the nucleus. Compartmentalization is readily detected in Hi-C maps by the presence of a checkerboard-like pattern. This pattern emerges because compartmental domains of similar chromatin status within the same and on different chromosomes interact with one another preferentially, producing an off-diagonal checkering effect on contact frequency. Underscoring the independence of these two organizing processes, depletion of cohesin eliminates TAD-associated patterns in Hi-C maps (Rao et al., 2017; Schwarzer et al., 2017; Wutz et al., 2017) and depletion of CTCF erases TAD boundaries (Nora et al., 2017), and in both cases compartmentalization patterns remain intact.

The details of compartmentalization in the nucleus remain unclear. The possible mechanistic contributions to observed compartmentalization are thought to include (i) microphase separation driven by specific biochemical affinities between chromatin types, most likely dictated by their epigenetic state (histone modifications and recruited factors) and (ii) the association of certain regions of the genome with nuclear bodies (speckles, nucleolus) and tethering to the nuclear periphery (lamina). Importantly, because of the slow timescale of global compartmental organization in interphase, it can also be influenced by the configurations of chromosomes upon exit from mitosis (Abramo et al., 2019; Luperchio et al., 2018; Zhang et al., 2019). Loop extrusion and compartmentalization are driven by distinct mechanisms that operate independently but act simultaneously and therefore can interfere with each other. For example, simulations have shown that the ATP-driven process of loop extrusion can locally counteract the demixing effect of affinity-driven compartmentalization (Nuebler et al., 2018).

Previous simulations of chromosome compartmentalization in inverted nuclei strongly indicated that attractions between heterochromatic loci are the major force driving compartmentalization (Falk et al., 2019). Heterochromatin is usually categorized into two types. Facultative heterochromatin is associated with Polycomb repressive complexes 1 and 2 and enriched in H3K27me3 (Penagos-Puig & Furlan-Magaril, 2020). It is considered to form at regions containing genes that are silenced in a developmentally regulated manner. Constitutive heterochromatin is viewed as more static, is primarily associated with H3K9me3, and forms at centromeres, pericentromeric regions, and at telomeres (Janssen et al., 2018). However, H3K9me3-associated heterochromatin is also found to form large contiguous domains genome-wide that expand in number and size during differentiation from pluripotency (Becker et al., 2016). H3K9me3 heterochromatin is associated with HP1 proteins, of which three homologs exist in mammals: HP1α, HP1β, and HP1γ. HP1 proteins bind H3K9me3 (reviewed in (Allshire & Madhani, 2018)) and can self-oligomerize and recruit H3K9 methyltransferases potentially contributing to heterochromatin compaction (Canzio et al., 2011; Machida et al., 2018), spread (Al-Sady et al., 2013; Müller et al., 2016), and phase separation (Larson et al., 2017; Sanulli et al., 2019; Strom et al., 2017).

Heterochromatin formation and maintenance is associated with DNA methylation and DNA methyltransferases. For example, megabase-long domains of reduced DNA methylation coincide with methylated H3K9 heterochromatin in various primary tissues, immortalized and cancer cell lines (Hansen et al., 2011; Salhab et al., 2018; Wen et al., 2009). Furthermore, in humans, the DNA methyltransferase DNMT1 physically associates with all three HP1 proteins and H3K9 methyltransferase G9a suggesting an interplay between DNA and histone methylation (Estève et al., 2006; Smallwood et al., 2007). CTCF-DNA binding has also been demonstrated to be DNA methylation-sensitive through CpG methylation of the core binding motif (Bell & Felsenfeld, 2000; Hark et al., 2000; Phillips & Corces, 2009; Wang et al., 2012). Overall, the regulatory relationships between the DNA methylome, CTCF, and heterochromatin are likely critical for cell-type specification and are still poorly understood.

Early analyses of Hi-C maps were able to subdivide mammalian genomes on the basis of long-range contact frequencies into two groups or “compartments” termed A and B, broadly correlating with transcriptionally active and inactive chromatin (Imakaev et al., 2012; Lieberman-Aiden et al., 2009). Higher resolution Hi-C data has shown that this binary organization is too simplistic. Until recently, most discussion of finer classifications of compartmental structure have largely focused on a single deeply sequenced immortalized lymphoid cell line, GM12878 (Rao et al., 2014). However, profiles of long-range contact frequency are not locus-intrinsic properties. Since the Hi-C profile of a single locus depends on the chromatin state of the remainder of the genome, long-range patterns can be difficult to generalize and compare across cell types with divergent chromatin landscapes. Conversely, even when congruences are found where a group of loci share similar interaction profiles in each of two different cell types, there is no guarantee that the underlying chromatin states are identical.

Here, we report a detailed investigation of nuclear compartmentalization motivated by the prominent compartmentalization of heterochromatin in HCT116 colon cancer cells. Using unsupervised methods and integrative analysis, we are able to distinguish three inactive chromatin states on the basis of long range contact frequencies, including a previously uncharacterized interaction profile marked by H3K9me2 and the histone variant H2A.Z that displays a neutral interaction preference. We find a strong compartmentalization signature for heterochromatin marked by megabase-long continuous stretches of H3K9me3, HP1α and HP1β that (i) lacks loop extrusion barrier-associated patterns in Hi-C, (ii) is abundant throughout the genome in HCT116, yet (iii) is present in all cell types we investigated, with the lowest in embryonic stem cells and GM12878. We demonstrate that this heterochromatin is lost under the influence of both acute DNA methylation inhibition and DNA methyltransferase knockout in HCT116, dramatically altering genome compartmentalization but not replication timing. Finally, we reveal an interplay between heterochromatin and loop extrusion: (i) H3K9me3-HP1α/β heterochromatin is permissive to loop extrusion by cohesin but is depleted for CTCF extrusion barriers, (ii) regions that lose H3K9me3-HP1α/β heterochromatin undergo widespread reactivation of CTCF sites, and that (iii) this outcome is due not to canonical demethylation of the CTCF motif but likely an intrinsic resistance of heterochromatin to CTCF binding. Taken together, these results highlight diversity and plasticity in heterochromatin, and its influence on the two major chromosome-organizing processes in interphase.

## Results

### Spectral decomposition of Hi-C identifies distinct interaction profiles

There is evidence that some cell lines or cell types may have unique nuclear compartmentalization features. For instance, HCT116 colon cancer cells appear to harbor compartmentalization signatures in Hi-C that are distinct from other cell lines (Nichols & Corces, 2021; Xiong & Ma, 2019). Furthermore, using Protect-seq, which measures peripheral heterochromatin, we have previously shown that in HCT116, H3K27me3 and H3K9me3 distinguish between two structurally distinct types of heterochromatin (Spracklin & Pradhan, 2020). Therefore, we sought to identify groups of loci through similarities in long-range interaction profiles (i.e. rows of Hi-C maps) and to understand the relationship of these groups to the chromatin landscape in HCT116. (**Figure 1A**). Our method for characterizing interaction profiles leverages the information from *trans* interactions as in (Rao et al., 2014) but introduces an initial dimensionality reduction step similar to (Lucic et al., 2019). Rather than cluster Hi-C contact matrices directly, we replace the contact frequency data of individual loci with a dimensionally reduced representation (i.e. leading eigenvectors, see **Methods**). We apply *k*-means clustering to the eigenvectors to identify clusters that represent groups of loci having similar interaction profiles. The rationale for this approach is that it is computationally efficient, dampens noise, and highlights cluster structure in the data (Brand & Huang, 2003). Unlike previous methods it does not require harmonizing cluster labels between different data subsets. Extracting the leading eigenvectors from Hi-C maps also facilitates the projection and embedding of genomic loci to allow investigation of the continuous structure of the interaction profile manifold, in which each point corresponds to a 50-kb genomic bin (**Figure 1B**).

**Figure 1:**
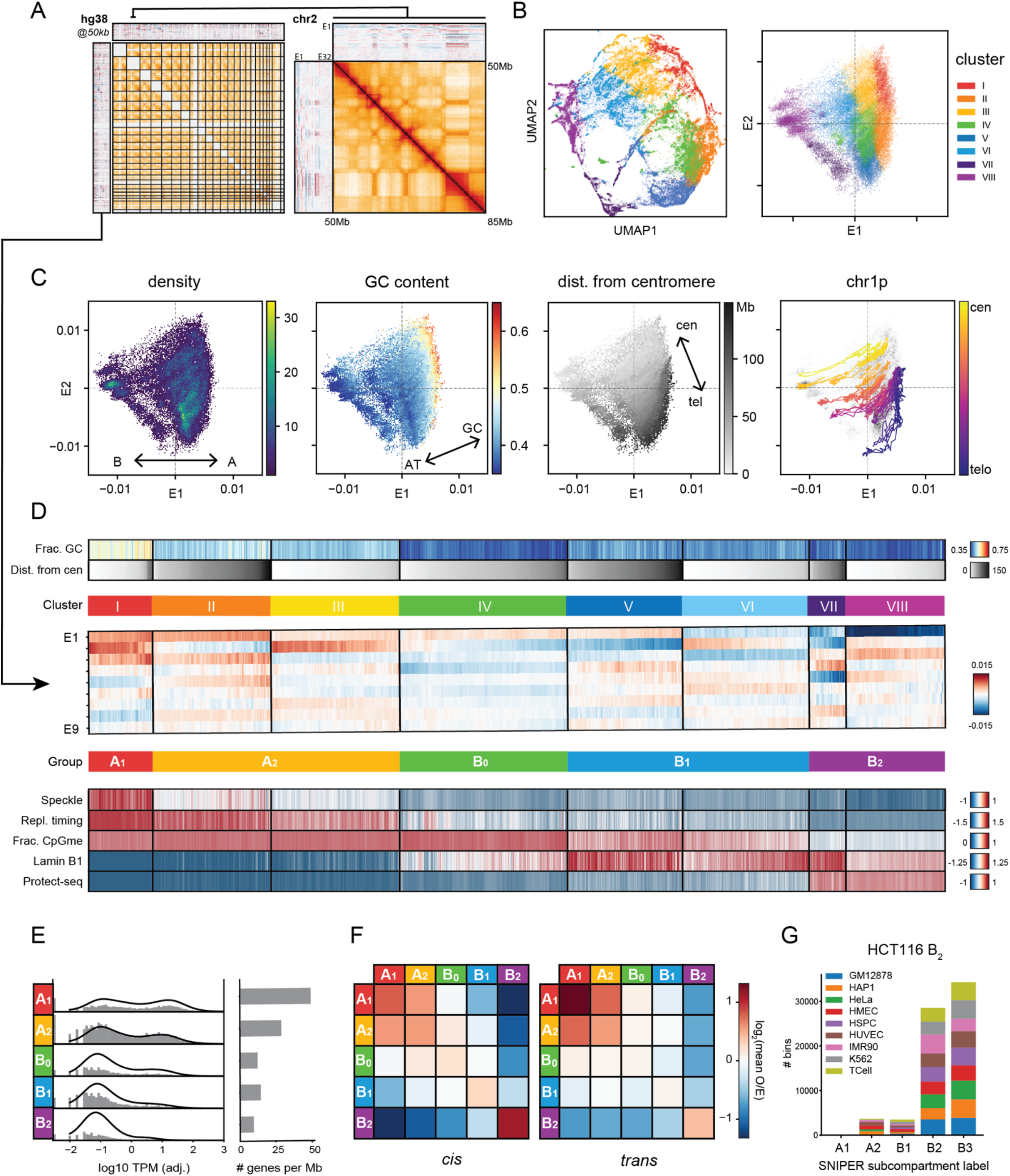
Spectral decomposition of *trans* Hi-C data identifies distinct interaction profiles. (A) Left, Hi-C contact map of HCT116 with masked intrachromosomal zones and heatmaps of leading *trans* eigenvectors displayed in both the top and left margins; right, blowup of a 35-Mb intrachromosomal region of chromosome 2; (B) Scatter plots of all 50kb genomic bins projected onto (left) a UMAP embedding of the nine leading eigenvectors (E1-E9) and (right) the E1-E2 subspace, colored by cluster identity using *k*-means on E1-E9 (*k*=8). (C) E1 vs. E2 scatter plots of 50kb genomic bins colored by (from left to right) point density, GC content and distance from the centromere. The fourth plot contains the trajectory of a single chromosome arm (chr1p) traced from centromere to telomere through the E1-E2 subspace (colored) on top of the point density in grayscale. (D) Heatmaps of mean signal intensity of functional genomics features and assays (rows) for each 50kb genomic bin (column), grouped into spectral clusters as in (B). The displayed features from top to bottom are GC content, distance from centromere, TSA-seq for Speckle-associated protein SON, two-stage Repli-seq (Early/Late), fraction of methylated CpGs derived from WGBS, Lamin B1 DamID-seq, and Protect-seq. The clusters (I-VIII), labeled with colored bars, are displayed in ascending order of Protect-seq signal and the bins within each cluster are sorted by distance from the centromere. Three pairs of clusters are combined based on feature similarity, giving a total of five interaction profile groups: A_1_, A_2_, B_0_, B_1_, B_2_, indicated in the second row of colored bars. A heatmap of the E1-E9 tracks used for clustering is shown in between. (E) Left, total RNA expression per interaction profile group represented as a log-scaled histogram of Transcripts Per Million (TPM) with group-normalized KDE plots as solid lines. Right, average number of genes per megabase in each interaction profile group. (F) Heatmap displaying the pairwise mean observed/expected contact frequency between interaction profile groups at 50kb in *cis* (left) and *trans* (right). (G) Distribution of SNIPER subcompartment label assignments (Xiong and Ma 2019) of HCT116 B*_2_* loci across various cell types.

In contrast to the discrete compartment paradigm, we observe that the manifold does not form dense, strongly-separated clusters as evidenced by the relatively continuous UMAP embedding of the leading eigenvectors (**Figure 1B**, see **Methods**). Furthermore, projecting loci onto the two leading eigenvectors (E1 and E2), we notice that GC content and genomic distance from centromere of individual loci vary along almost perpendicular components in the projection (**Figure 1C**). A similar pattern is observed in other cell types besides HCT116 suggesting it represents a conserved feature (**S. Figure 1**). The alignment of GC content to E1 is well known, but the exact relationship differs across cell types (Imakaev et al., 2012). The positional component correlating strongly with E2 reflects the observation that pairs of centromere-proximal and centromere-distal regions show mildly elevated contact frequency throughout the genome (**Figure 1C**) (Imakaev et al., 2012). This may be due to known enrichment of interactions between telomeres and/or between centromeres, or a relationship between chromosomal and nuclear landmarks during interphase. As a result, we expected that the clustering of interaction profiles using *trans* Hi-C data would be influenced by chromosomal position independently of chromatin state. To test this idea, we examined subcompartment calls from GM12878 (Rao et al., 2014). Indeed, the inactive interaction profiles B2 and B3 in GM12878 appear to differ positionally along the E2 axis (**S.Figure 2**). Similarly, in HCT116 cells we observe that several pairs of clusters with similar E1 ranges separate along the E2 axis (**Figure 1B**). These data suggest that the clustering of some interaction profiles will be influenced by positional correlates that are likely unrelated to biological function.

We find that the data can be sensibly partitioned into eight clusters (see **Methods**, **S.Figure 1**). To exclude the influence of genomic position, we next examined functional genomic assays (e.g. replication timing, lamin association, nuclear speckle association, and nuclease resistance) including publicly available data (**Supplemental Table 1**) (Dekker et al., 2017; ENCODE Project Consortium, 2012). We observe that several clusters were enriched for distinct sets of genomic features (**Figure 1D**). We therefore consolidated centromere-proximal and distal pairs of clusters with similar functional profiles for a total of five groups (described in detail below). For simplicity, we have chosen a similar naming system to those used in clustering GM12878 *trans* interaction profiles, but below we discuss what correspondences can be made.

We identify two transcriptionally active groups, consistent with previous reports (Rao et al., 2014). Cluster I has the strongest self-interaction preference in *trans*, is enriched for the nuclear speckle marker SON, and displays the greatest amount of transcriptional activity (**Figure 1D**, **1E**, **1F**). Its loci have a high degree of overlap with the A1 interaction profile identified in GM12878 cells and thus we termed this profile A_1_ (**S.Figure 1**). In GM12878, A2 has been described in more generic terms as domains with weak transcriptional activity. Thus, clusters II and III which display weak transcriptional activity and separate along the E2 axis (cen/telo) are grouped and classified as A_2_ (**Figure 1B, 1E**). Interestingly, the A_2_ group interacts with the A_1_ group (heterotypic) at least as strongly as it does with itself (homotypic) (**Figure 1F**). This lack of self-preference suggests that A_2_ is unlikely to represent well-defined spatial compartments and that it does not spatially separate from A_1_.

The five remaining clusters all display low transcriptional activity and gene density and thus likely constitute inactive chromatin domains (**Figure 1E)**. Clusters VII and VIII are both enriched in Protect-seq signal, are late-replicating, display the lowest CpG methylation frequency and have the strongest preference for homotypic contacts in *cis* (**Figure 1D**, **1F**). The majority of loci in these clusters are assigned subcompartment labels B2 and B3 in GM12878 and are consistently assigned labels B2/B3 across different cell types based on SNIPER, a supervised model that generalizes the GM12878 labels to other cell types (**Figure 1G** and **S.Figure 1**). However, as will be shown below, in HCT116 these loci harbor the mark H3K9me3, which is most similar to the GM12878 B4 subcompartment (previously found uniquely on chromosome 19 (Rao et al., 2014), although we detect an expanded B4 cluster in GM12878 using our method - **S.Figure 2**). For simplicity, we assign clusters VII and VIII the name B_2_ due to the high degree of overlap, but emphasize that no functional correspondence to the B2 or B3 GM12878 labels is implied by the name. Clusters V and VI are both enriched in Lamin B1, are late replicating, and intermediate CpG methylation consistent with the B1 subcompartment label and thus we term this group B_1_ (**Figure 1D**). Compared to B_2_ loci, loci in B_1_ have more diverse subcompartment labels in different cell types, which is consistent with facultative heterochromatin (**S.Figure 1**).

Interestingly, we identified a novel type of interaction profile (cluster IV) with no equivalent in GM12878, whose loci share hallmarks of inactive chromatin along with B_1_ and B_2_ such as low GC content, late replication timing, and association with the lamina (**Figure 1D**). Despite low GC content, it exhibits high CpG methylation frequencies and unlike B_2_, it does not exhibit Protect-seq enrichment (**Figure 1D**). This cluster has a distinct 3D interaction profile, showing only modest preference for homotypic contacts (**Figure 1F**), suggesting these do not form well defined spatial subnuclear compartments. However, cluster IV regions do form large continuous domains, present on many chromosomes (**S.Figure 1**). And importantly, when cluster IV loci are compared to subcompartment labels in other cell types it appears to be either weakly transcriptionally active (A2) or silent (B3) (**S.Figure 1**), suggesting this cluster could represent a ‘poised heterochromatin’ that transitions between active and inactive chromatin in different cell types. As such, we have termed this cluster B_0_ and speculate that it could indicate a potentially bivalent underlying chromatin state (see below).

In summary, our spectral clustering method is a simple and scalable alternative to identify signature interaction profiles for classifying genomic loci that is comparable to prior methods. In addition, we conclude that, by discounting positional effects, HCT116 cells have five distinguishable groups of loci identifiable on the basis of long-range contact frequencies, including a previously uncharacterized inactive group, B_0_ (**Supplemental Table 2**). Moreover, these data highlight the potential pitfalls of extrapolating interaction profile assignments between distant cell types. And importantly, not all interaction profiles imply the existence of phase-separated subnuclear compartments. Therefore, from here we refer to our classification labels as interaction profile groups rather than (sub)-compartments. We subsequently focused on distinguishing and assigning biological function to the interaction profile groups.

### 3D interaction profiles discern three types of inactive chromatin in HCT116

To understand the chromatin composition of the five interaction profile groups described above, we examined histone modifications, histone variants and related factors (**Figure 2A**). Interaction profile groups A_1_ and A_2_ are enriched for histone modifications associated with active transcription, similar to what has been found for other cell types (**Figure 2B**) (Rao et al., 2014; Xiong & Ma, 2019). Consistent with B_1_ being facultative heterochromatin, these loci are predominantly enriched for H3K27me3, with a mild enrichment in H3K9me2 (**Figure 2B** and **2C**). B_0_ also displays a subtle enrichment in H3K9me2 (**Figure 2B** and **2C**). With that said, H3K9me2 is a broad histone mark and datasets from different groups show subtle differences in the degree of enrichment in B_1_ domains and thus should be interpreted with caution. Loci with the B_2_ interaction profile are marked with H3K9me3, HP1α and HP1β, consistent with these loci being in a constitutive heterochromatic state (**Figure 2B** and **2C**). To further investigate the inactive chromatin landscape of B_0_, B_1_, and B_2_, we applied a multivariate Hidden Markov Model (ChromHMM) to segment the genome into states based on combinations of chromatin features (see **Methods**). Mapping those chromatin states to our interaction profile groups, we observe that B_0_ is almost entirely composed of the HMM state that emits H3K9me2 without H3K27me3 (**S.Figure 3**). Finally, when the E1-E2 projection of loci is colored by H3K27me3 or H3K9me3 an enrichment pattern spans the entire E2 axis, further validating the assumption that consolidated centromere/telomere-proximal clusters are functionally similar (**Figure 2D** and **S.Figure 3**).

**Figure 2:**
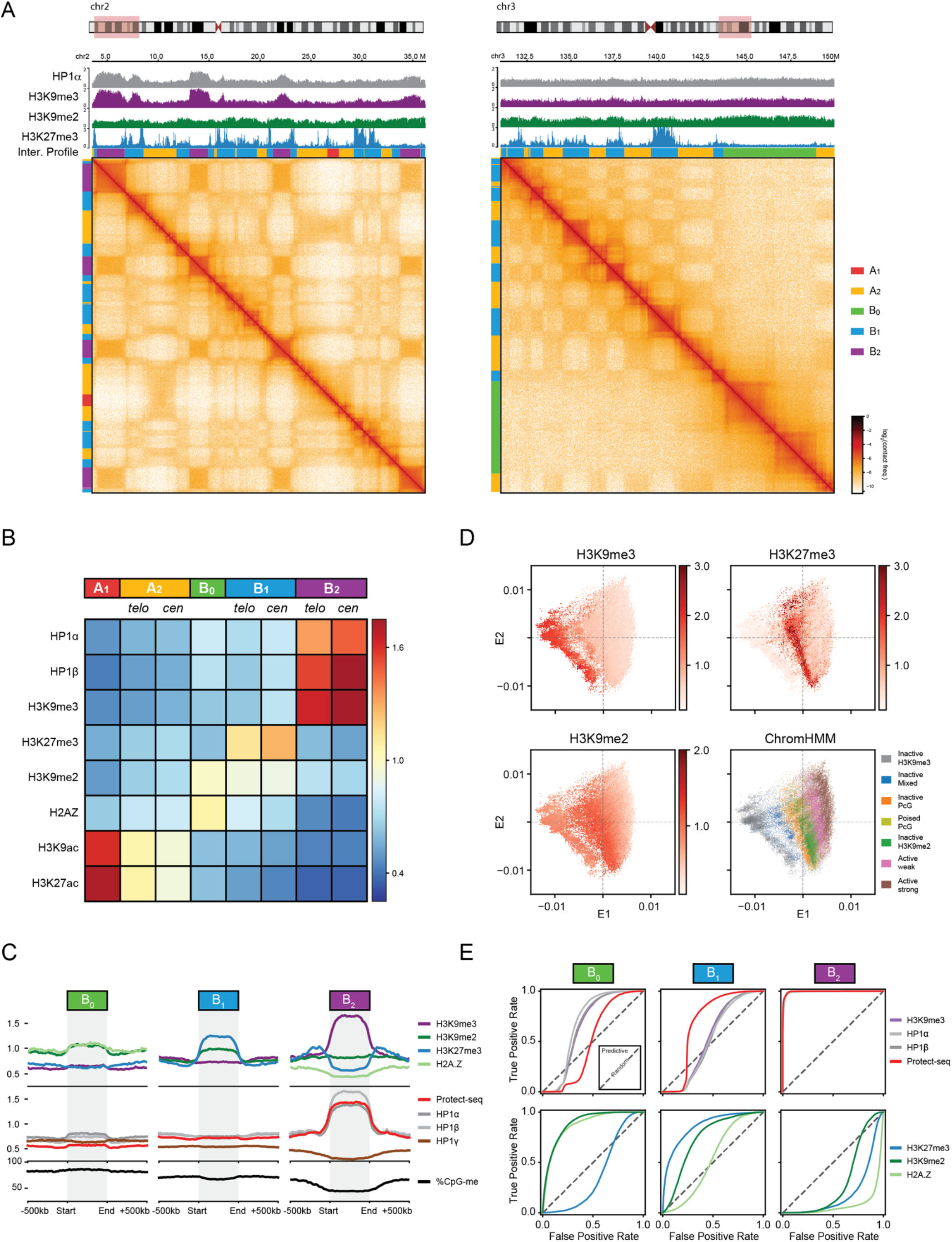
3D interaction profiles discern three types of inactive chromatin in HCT116. (A) Two example regions illustrating the contrasting interaction profiles of B_2_ domains (left, chr2:3.5-3.6M) and B_0_ domains (right, chr3:131-150M) against A_1_, A_2_, and B_1_ in *cis*. The interaction profile group labels are displayed as colored bars on the top and left margins (A_1_, red; A_2_, yellow; B_0_, green; B_1_, blue; B_2_, purple). Top, ChIP-seq tracks for HP1α, H3K9me2, H3K9me3, and H3K27me3. (B) Heatmap of mean fold-enrichment of ChIP-seq signal intensity for histone modifications and HP1α and HP1β proteins averaged over 50-kb bins in each interaction cluster (*k*=8). (C) Metaplots of B_0_, B_1_, B_2_ domains displaying signal enrichment for ChIP-seq (H3K27me3, H3K9me2, H3K9me3, H2A.Z, HP1α/β/γ), Protect-seq, and DNA methylation rescaled to 25 bins and flanked by ±500kb (D) E1-E2 scatter plots of 50kb bins colored by ChIP-seq signal enrichment (H3K27me3, H3K9me2, H3K9me3) and ChromHMM state annotation. (E) Receiver operating characteristic (ROC) curves assessing the prediction performance of individual 50kb-aggregated functional tracks (ChIP-seq, Protect-seq) when treated as binary classifiers for B_0_, B_1_ or B_2_ loci. The discrimination threshold parameter in each case is a simple global binarization threshold on the signal track.

**Figure 3:**
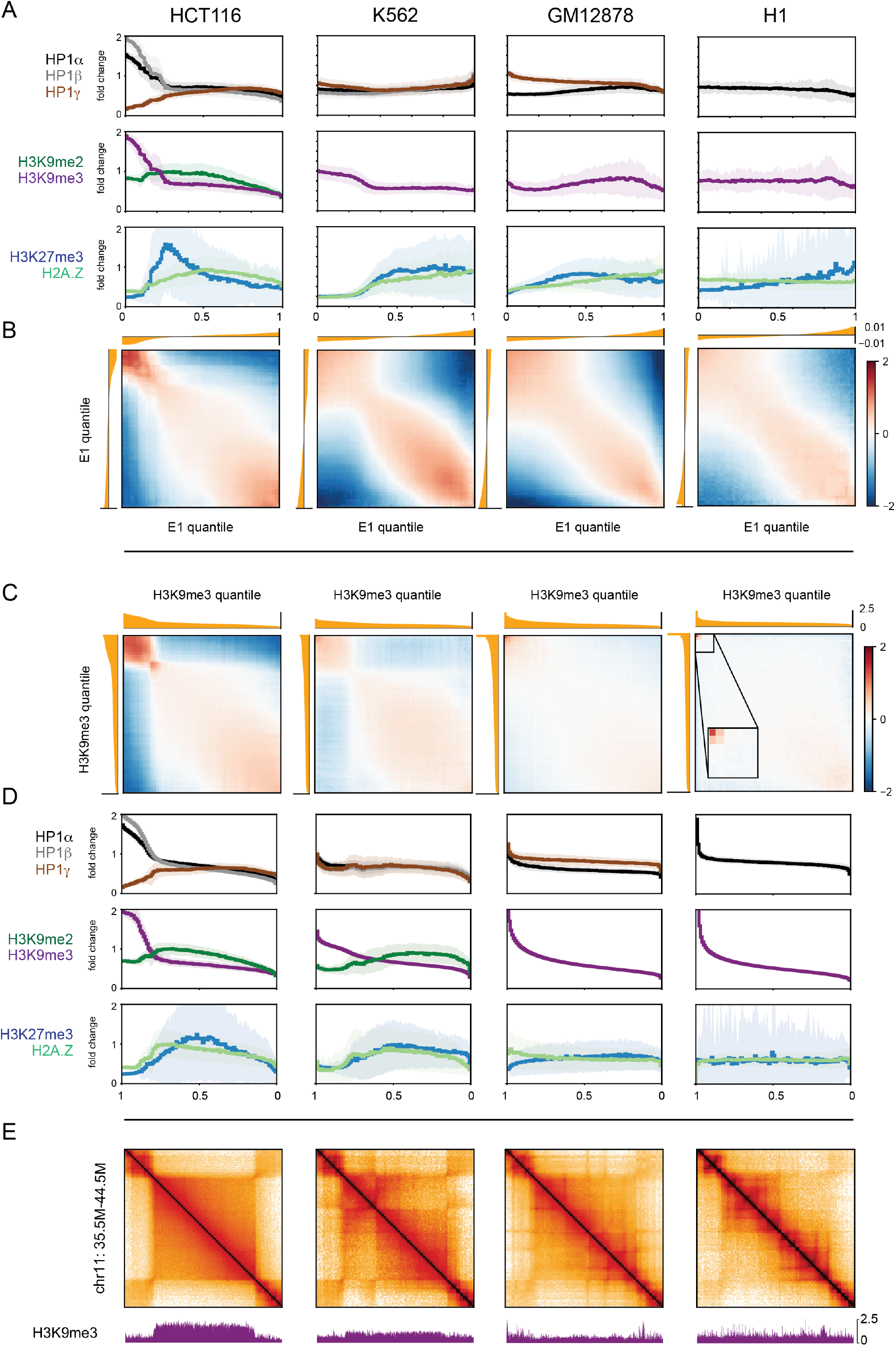
Comparative analysis indicates a wide prevalence range of chromatin marked by H3K9me3, HP1α and HP1β and strong homotypic interaction preference. Comparative analysis of genome organization and heterochromatic marks across HCT116, K562, GM12878 and H1-hESC. (A) Histograms of ChIP-seq signal for repressive histone marks, HP1 proteins and H2A.Z grouped by eigenvector (E1) percentile and displayed in ascending order of E1 rank. Solid lines display the mean over the 50kb bins within each percentile and include a standard deviation envelope. (B) Bivariate histograms of observed/expected contact frequency (a.k.a. saddle plots) based on E1 percentiles and aligned with the univariate ChIP-seq histograms above in (A). (C) Bivariate histograms similar to (B) but based on percentiles of H3K9me3 signal, displayed in descending order of H3K9me3 rank. (D) Histograms of ChIP-seq signal similar to (A) but based on percentiles of H3K9me3 signal, aligned with the bivariate histograms in (C). (E) Hi-C maps of a region containing a B_2_ domain in HCT116 (chr11:34.5-44.5M) and corresponding H3K9me3 signal below.

Curiously, in addition to H3K9me2, B_0_ also has a mild enrichment for the histone variant H2A.Z (**Figure 2B** and **2C**). In humans, hypoacetylated H2A.Z has been reported to co-exist with H3K9me2 in broad lamina-associated chromatin domains suggesting that interaction profile group B_0_ could correspond to a similar type of chromatin (Giaimo et al., 2019; Hardy et al., 2009; Kafer et al., 2020). Moreover, B_0_-like domains that display neutral interaction profiles in Hi-C and broad H3K9me2/H2A.Z chromatin modifications are detected in other cell types, e.g. human umbilical vein cells (HUVEC) suggesting this interaction profile is unlikely an artifact of HCT116 cells (data not shown). The biological function of the B_0_ interaction profile remains unclear; however, the presence of H3K9me2 and H2A.Z further supports our speculation that B_0_ domains are a poised state, i.e. an inactive chromatin state that maintains the potential to have an euchromatic or heterochromatic profile in further differentiated cells.

Our A_1_ and B_2_ assignments (7.5% and 15.9% of the genome, respectively, excluding unmappable and translocated regions) exhibit the closest correspondence to known euchromatic and heterochromatic chromatin states, respectively. This can be observed using ROC curves generated by using thresholded 50-kb binned signal tracks as binary classifiers for individual group assignments (**Figure 2E** and **S.Figure 3**). The A_1_ label is predicted by the nuclear speckle marker SON with an AUC of 0.986, and the B_2_ label is predicted by each of H3K9me3, HP1α, HP1β and Protect-seq with AUC > 0.992. These close correspondences, coupled with A_1_ and B_2_ being the most self-interacting groups, suggest that homotypic affinity could be drivers of A_1_ and B_2_ compartmentalization. Other interaction profile groups are less well predicted by any single chromatin modification, even though a particular histone modification may be globally enriched. We note that in general, even if chromatin state underlies the driving forces for the spatial segregation of chromatin in the nucleus, interaction profiles are not necessarily perfect proxies for chromatin state since other processes may exert influence on or reduce the contrast of long-range interaction patterns in contact frequency maps (e.g. loop extrusion). Indeed, when we identify interaction profile groups in Hi-C data from HCT116 cells in which the cohesin subunit RAD21 is depleted, we observe a slight increase in correspondence to ChromHMM labels (Adjusted Rand Index: HCT116 = 0.31, HCT116-RAD21 = 0.35) (**S.Figure 3**). This is consistent with loop extrusion interfering with innate compartmentalization preferences (Schwarzer et al., 2017).

Finally, we note that HMM analysis reveals an additional mixed state that emits a combination of H3K9me3-HP1α/β (similar to B_2_) and H3K9me2 (similar to B_0_) (**S.Figure 3**). Upon closer inspection, we see that some of these regions correspond to domains having continuous ChIP-seq and DNA methylation profiles with fractional heights relative to their neighbors (**S.Figure 4**). A parsimonious interpretation for this observation is population heterogeneity or allelic imbalance. Moreover, the long-range Hi-C profiles of such regions also appear to be a superposition of B_0_ and B_2_ (**S.Figure 4**). These data are reminiscent of breast cancer cells which display allelic imbalance with one allele being enriched for repressive histone modifications (H3K9me3 or H3K27me3) and the other for DNA methylation (Hon et al., 2012).

In summary, we discern five functionally distinguishable groups of similarly interacting loci in HCT116 cells with A_1_ and B_2_ being the most homotypic or self-interacting and the most stable across cell types. Our cluster analysis also suggests at least three types of inactive chromatin are distinguishable by long-range contact frequencies in HCT116: B_0_ (H2A.Z and H3K9me2 but H3K27me3-depleted), B_1_ (H3K9me2 and H3K27me3-enriched) and B_2_ (H3K9me3, HP1α and HP1β). Notably, none of these types appear to share an epigenetic similarity with the B2/B3 subcompartments described in GM12878 (**S.Figure 2**). These results therefore hint at a greater diversity of inactive chromatin types, within and between cell types, than broadly attested.

### Chromatin marked by H3K9me3, HP1α and HP1β displays a conserved homotypic interaction preference indicative of self-affinity

Our data suggests that B_2_ domains are enriched for H3K9me3, HP1α and HP1β and have strong homotypic interaction preferences. Next, we asked whether this observation in HCT116 is a conserved feature in other cell types. To explore this conservation, we examined HCT116, K562, GM12878, H1-hESC cell lines for which we could obtain high quality Hi-C data as well as ChIP-seq for HP1 proteins and repressive histone marks. First, we examined ChIP-seq enrichments of H3K9me2/3, HP1α/β/γ, H3K27me3, H2A.Z and binned them into quantiles according to E1 value (**Figure 3A**). These data confirm our previous observation that negative E1 quantiles (which correspond to the classical B compartment) are enriched for H3K9me3, HP1α and HP1β in HCT116. K562 cells also displayed an enrichment in H3K9me3 in negative E1 quantiles, albeit weaker (**Figure 3A** and **S.Figure 5**). In GM12878 cells we observed lower abundance of H3K9me3, and H3K9me3 was also found in active regions with positive E1 values. Human embryonic stem cells (H1) have an even lower abundance of H3K9me3 (**Figure 3A**), consistent with microscopy data suggesting H1 lacks punctate constitutive heterochromatin (Mattout et al., 2015; Ugarte et al., 2015).

Next, we sought to understand whether the presence of H3K9me3, HP1α, and HP1β was correlated with preferential homotypic interactions, thus potentially forming sub-nuclear compartments. To do this, we profiled *cis* contact frequency between pairs of loci ranked by their E1 eigenvector status and compared this to *cis* contact frequency between pairs of loci ranked by H3K9me3 enrichment. We find that loci with similar E1 status tend to interact with each other, as expected (**Figure 3B**). Interestingly, loci that display high levels of H3K9me3 also show particularly high contact frequencies with each other (**Figure 3C** and **S.Figure 5**). This phenomenon is observed in all cell types even though GM12878 and H1 have a much lower abundance of H3K9me3 loci than HCT116. Loci in the highest H3K9me3 quantiles also show elevated HP1α in all cell types as well as HP1β where data was available (**Figure 3D**). We conclude that the presence of H3K9me3 along with HP1α and HP1β is correlated with elevated homotypic contact frequency across cell types regardless of genomic abundance. Interestingly, in GM12878 and K562 we also observe a co-enrichment of HP1γ with H3K9me3, while HP1γ is anticorrelated with H3K9me3/HP1α in HCT116 (data for H1 was unavailable).

HCT116 cells have large ungapped H3K9me3 (B_2_) domains up to several megabases in length (**Figure 3E** and **S.Figure 6**). Therefore, we asked if broad H3K9me3-enriched domains are a conserved feature across six cell types. Taking the largest domains in each cell type ranked by size, we observe that K562 and fibroblasts (HFFc6, IMR90) also exhibit large domains (**S.Figure 6**). In GM12878 and H1 cells we observed shorter domains compared to HCT116 and K562 (**S.Figure 6**). Yet even among the few domains in H1 cells displaying H3K9me3 and HP1α, we observe a tendency to self-interact (**S.Figure 6**).

It is noteworthy that, in contrast to *cis* contact frequency, *trans* contact frequency between H3K9me3-containing loci is not generally elevated across cell types (**S.Figure 5**). These data argue that chromosomal territoriality and/or association with nuclear landmarks (e.g. lamina) can limit the extent of interchromosomal contacts between H3K9me3 loci. Finally, the fact that loci with similar E1 value show preferred interactions with each other across the full range of E1 values, indicates that other factors besides H3K9me3-HP1 can also mediate such interactions. For instance, in K562, GM12878 and H1 cells loci with low/negative E1 values still prefer to interact with other loci with similar E1 values even though in these cells most of these loci do not display strong H3K9me3-HP1 enrichment (**Figure 3B**).

Taken together, these data suggest that the constitutive heterochromatin marks, H3K9me3 and HP1, define a homotypically interacting chromatin state, but that the prevalence and distribution of this chromatin state varies substantially across cell types. The exact combination of HP1 homologs and/or post-translational modifications may also exert an influence (Lomberk et al., 2006). Interestingly, the large, continuous plateaus of H3K9me3-HP1α/β enrichment in HCT116 correlate with large, smooth B_2_ domains in Hi-C maps that lack loop extrusion-associated features such as TADs, dots, and stripes (**Figure 3E** and **S.Figure 6**). From our cell type comparisons, we conclude that the strength of heterochromatic homotypic interactions and their contribution to overall genome compartmentalization as observed in contact frequency maps depends on the prevalence, distribution and population-level saturation of H3K9me3 and HP1 proteins along the genome. Consistent with the putative role of HP1 proteins as cross-linkers, it is likely that H3K9me3-HP1α/β compartmentalization is driven directly by HP1-mediated affinity.

### Chromatin is ubiquitously permissive to loop extrusion but H3K9me3-HP1α/β heterochromatin lacks extrusion barriers

Besides compartmentalization, another major organizing mechanism in the nucleus is loop extrusion. The signature patterns of TADs, dots and stripes, arise as a combination of two effects: (1) extrusion of loops by cohesin, and (2) CTCF acting as an extrusion barrier. Such patterns are readily visible in Hi-C contact maps and appear most abundantly in A-type domains, and rarely within B_2_ domains in HCT116 cells. We therefore set out to determine whether a lack of extrusion or a lack of CTCF barriers (or both) is responsible for the non-appearance of loop extrusion features in regions characterized as B_2_.

First, we examined B_2_ domains in cells with normal CTCF barriers but without extruding loops (*i.e.* Rad21 depleted HCT116 RAD21-AID cells) (Rao et al., 2017). We looked at the decay of contact probability with genomic separation, *P*(*s*), which is indicative of the underlying polymeric folding of the region. We found that *P*(*s*) was affected by depletion of cohesin in all interaction profile groups, including B_2_ domains, leading to the disappearance of the characteristic extrusion “shoulder” in *P(s)* (**Figure 4A**) (Fudenberg et al., 2017). In the loop extrusion model, the size and density of extruded loops affects the shape of the *P*(*s*) curve (Flyamer et al., 2017; Gassler et al., 2017). A local maximum in the derivative at genomic distances of around 100-200 kb reflects the average loop size, which appears to be somewhat larger for B_2_ domains. We find that the shape of the *P(s)* derivatives suggest that A_1_ and A_2_ domains have more loops per kb than B_2_ (**Figure 4A**). These data support that all domain types characterized by interaction profiles accommodate extrusion of chromatin loops but with varying loop size and density, with B_2_ domains having relatively large loops and low loop density.

**Figure 4:**
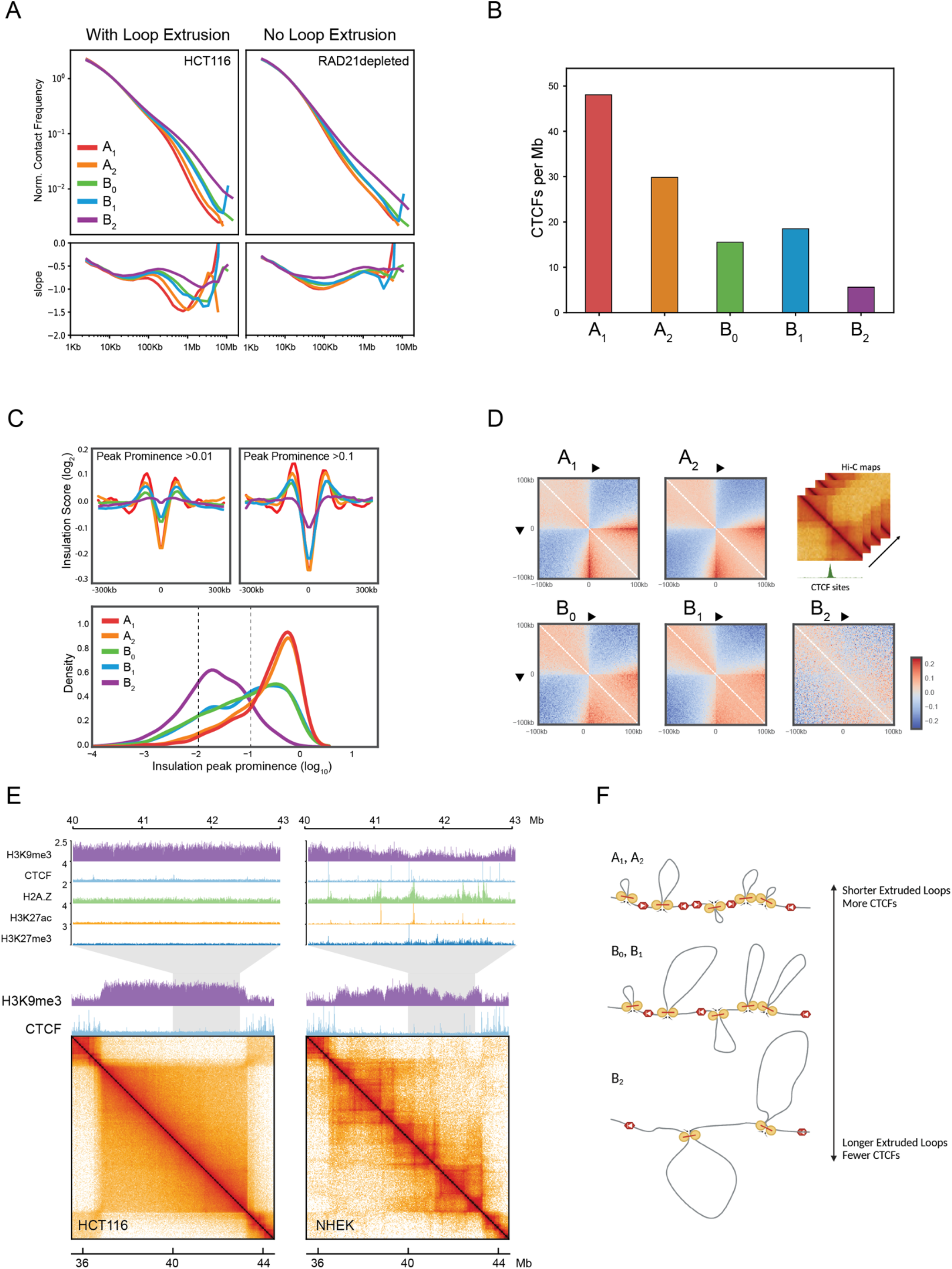
All interaction profile groups in HCT116 are permissive to loop extrusion but B_2_ domains lack extrusion barriers. (A) Top, *P*(*s*) curves, or interaction frequency as a function of genomic distance, for HCT116 and cohesin-depleted HCT116 RAD21-AID Hi-C, restricted to A_1_ (red), A_2_ (orange), B_0_ (green), B_1_ (violet), B_2_ (blue), normalized to unity at *s* = 10kb. Bottom, derivative of *P*(*s*) indicating average sizes of extruded loops regardless of appearance of dots and stripes in Hi-C data. (B) Mean number of CTCF peaks per megabase in each interaction profile group. (C) Top, average insulation score (log_2_) centered on 25-kb insulating loci (diamond size 100 kb) with peak prominence score greater than or equal to 0.01 (left) and 0.1 (right) per interaction profile group. Bottom, kernel density estimation plots of the insulation peak prominence (log_10_) distribution in each interaction profile group. Dashed lines indicate cutoffs for insulating loci used in panels above (>0.01 and >0.1 peak prominence). (D) Average observed/expected Hi-C maps around CTCF binding sites within each interaction profile group, centered at CTCF motifs oriented as indicated. Expected maps are calculated separately for each interaction profile group. (E) Contact frequency maps of a 9-Mb genomic region containing a B_2_ domain in HCT116 (chr11:35.5-44.5M) and the same region in normal human epidermal keratinocytes (NHEK) along with tracks for H3K9me3 and CTCF ChIP-seq. Top, zoom-ins of a 3-Mb subregion showing tracks for H3K9me3, CTCF, H2A.Z, H3K27ac and H3K27me3. (F) Model of extrusion barrier (CTCF) sparsity determining the average extruded loop size as reflected in the *P(s*) shoulder for each interaction profile group, with B_2_ domains having the fewest barriers and longest extruded loops.

Second, despite B_2_ domains appearing relatively featureless in HCT116 Hi-C maps, we find that extrusion-related stripes and dots (which disappear upon cohesin depletion) originating outside a domain can sometimes propagate through it, appearing along the periphery of the square (**S.Figure 7**). In the loop extrusion model, this would require the passage of extruded loops through the heterochromatic region, suggesting that featureless heterochromatic regions are traversable by cohesin. To test this hypothesis, we turned to polymer simulations of loop extrusion in a heterochromatic domain surrounded by tandem CTCF clusters. Stripes extending along the periphery of the B_2_ domains failed to appear when translocation of loop extrusion factors was blocked (**S.Figure 7**). These results argue that the loop extrusion machinery is able to traverse B_2_ domains.

Third, we find that the number of CTCF peaks is depleted in B_2_ domains compared to other interaction profile domains (**Figure 4B**). Moreover, the CTCF sites that are bound in B_2_ regions tend to show much lower enrichment (**S.Figure 7**). Concomitantly, we see fewer and weaker insulating loci in Hi-C at B_2_ domains (**Figure 4C**). Likewise, when we aggregate Hi-C data at CTCF-bound sites we find these sites form stripe-like features and local insulation (**Figure 4D**). For CTCF-bound sites in B_2_ domains these features are weak compared to these in other interaction profile groups, suggesting that weaker extrusion-associated features at sites in B_2_ domains can at least in part be explained by the low CTCF occupancy, and possibly by the lower levels of cohesin (**Figure 4A**). In contrast, when we examine CTCF in H1-hESC, which lack H3K9me3-HP1α/β chromatin in the same regions, we see similar occupancy levels across the genome (**S.Figure 7**). Altogether, our analysis argues that the low CTCF occupancy of B_2_ domains in HCT116 is not intrinsic to the DNA sequence, but rather that B_2_ domains in HCT116 are refractory to CTCF occupancy.

Finally, we also asked whether the lack of extrusion features in H3K9me3-HP1α/β regions are conserved across cell types. While we find it to generally be the case, we do find a subset of heterochromatic domains that have both broad H3K9me3 enrichment and also include extrusion-associated patterns in Hi-C (e.g. NHEK cells) (**Figure 4E**). We predicted that these subset of domains should have occupied CTCF binding sites at regions of low H3K9me3 saturation. Indeed, the visible TAD boundary loci have lower H3K9me3, are enriched for H2A.Z, and display narrow peaks for CTCF as well as marks such as H3K27ac and H3K27me3, suggesting that chromatin tends to be locally decompacted at these sites (**Figure 4E**). These data are reminiscent of ‘euchromatin islands’ previously described as small regions of CTCF occupancy embedded within large heterochromatin domains (Wen et al., 2012). The fact that dots and stripes can be detected in NHEK cells that cross domains enriched in H3K9me3 further supports that loop extrusion traverses heterochromatin.

Altogether, these data show that loop extrusion occurs within B_2_/H3K9me3-HP1α/β chromatin and that the absence of site-specific dots and stripes in Hi-C data is the result of low CTCF occupancy. Therefore, we put forth a model where the density of extrusion barriers differs across interaction profile domains resulting in different average extruded loop sizes (**Figure 4F**).

### DNA methyltransferase perturbation disrupts H3K9me3-HP1α/β heterochromatin and compartmentalization

Our data suggest that HP1/H3K9me3 heterochromatin is (i) abundant in HCT116, (ii) structurally unique, (iii) DNA-hypomethylated, (iv) depleted in CTCF binding, (v) and has strong homotypic interactions (B_2_ profile). Prompted by these observations, we sought out perturbation systems that disrupt H3K9me3-HP1α/β chromatin. First, the DNMT1 hypomorph and DNMT3b knockout cell line (termed DKO) (Rhee et al., 2002), is a DNA methylation-deficient line derived from HCT116 that has been reported to have defects in H3K9me3 (Lay et al. 2015) and HP1α/β deposition (Spracklin & Pradhan, 2020). To investigate acute effects of DNA methylation dysfunction on compartmentalization and chromatin state, we also examined HCT116 cells treated with 5-Azacytidine for 48h (hereafter referred to as 5Aza) (**Figure 5A**). In our hands, both the genetic and pharmacological conditions reduced DNA methylation compared to HCT116 cells as measured by LC-MS (**Figure 5B**).

**Figure 5:**
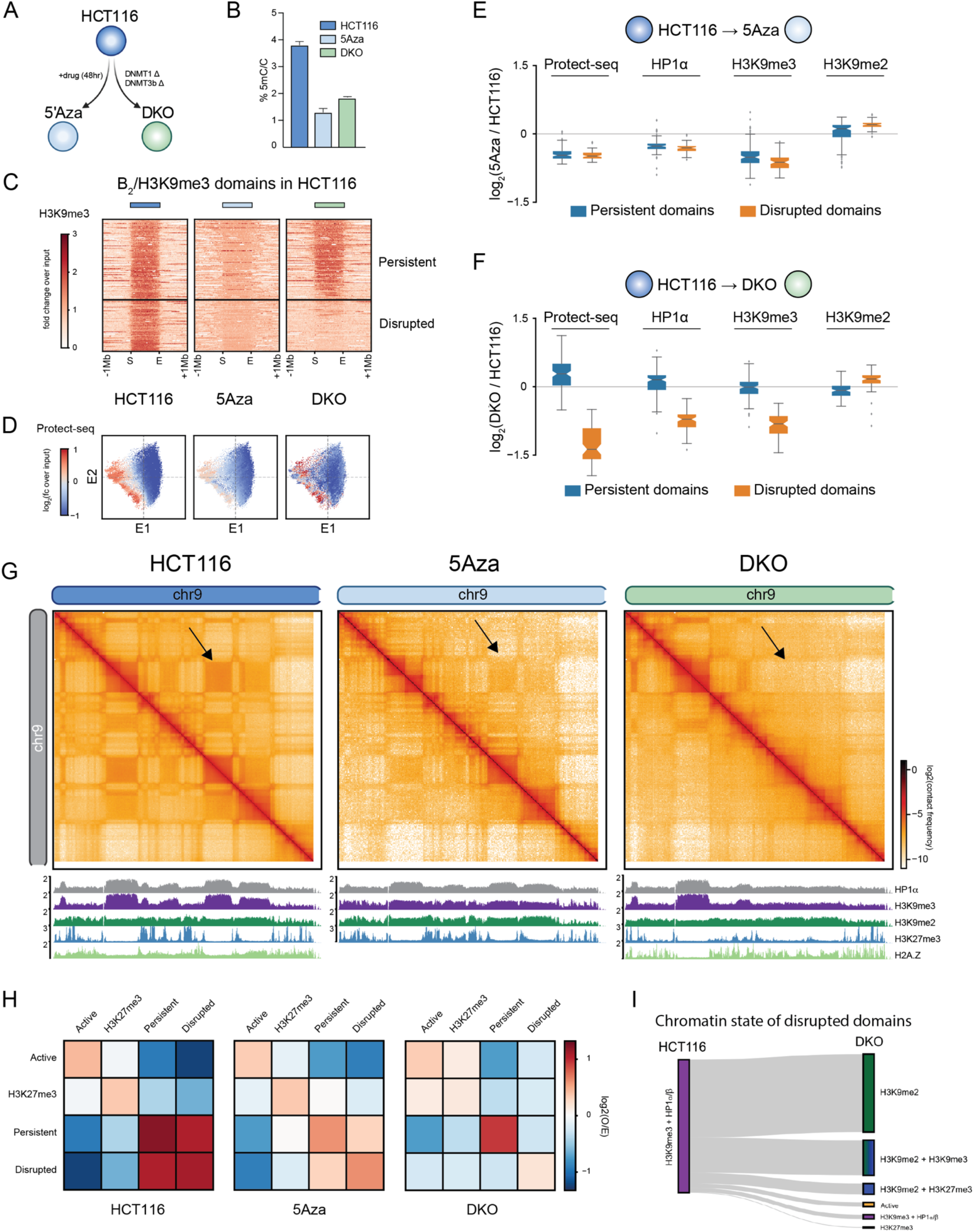
Inhibition or knockout of DNA methyltransferases disrupts H3K9me3-HP1α/β heterochromatin and compartmentalization. (A) Schematic of the DNA methylation perturbation system used in this study. (B) LC-MS quantification of 5-methylcytosine/total cytosine for HCT116 (left), HCT116 cells treated with 5-Azacytidine (5Aza) (middle), and DNMT1/DNMT3b knockout (DKO) (right) cells. (C) Stacked heatmaps of H3K9me3 ChIP-seq signal in HCT116 (left), 5Aza (middle) and DKO (right) centered at uniformly re-scaled B_2_ domains, sorted vertically by the intradomain H3K9me3 ratio between DKO and HCT116, and partitioned into two categories: persistent domains (top) and disrupted domains (bottom) in DKO. (D) Scatter plots of 50kb bins along E1 vs. E2 (HCT116 eigenvectors), colored by Protect-seq signal for HCT116 (left), 5Aza (middle), and DKO (right). (E) Boxplots quantifying the distribution of log_2_ ratios of mean domain signal between HCT116 and 5Aza in persistent (left) and disrupted (right) B_2_ domains. Signals shown are Protect-seq, HP1a, H3K9me3 and H3K9me2. (F) Same as (E) but between HCT116 and DKO. (G) Contact frequency maps of a 40-Mb genomic region (chr9:0-40M) in HCT116 (left), 5Aza (middle) and DKO (right) containing representative examples of persistent and disrupted domains. Below, ChIP-seq tracks for H3K27me3, H3K9me2, H3K9me3, HP1α, and H2A.Z. (H) Heatmap displaying the pairwise mean observed/expected contact frequency between active, H3K27me3, and H3K9me3 domains split into either disrupted or persistent labels in DKO based on ChromHMM states learned at 50kb. (I) Sankey plot of disrupted domains illustrating the chromatin transition from H3K9me3-HP1α/β in HCT116 cells to H3K9me2 and/or other repressive states based on ChromHMM in DKO cells.

As we have previously shown, in DKO cells a subset of domains are no longer detected by Protect-seq and no longer display HP1α and H3K9me3 binding, indicating that these domains are no longer in a closed heterochromatic state (**Figure 5C** and **S.Figure 8**) (Spracklin & Pradhan, 2020). This shows that not all B_2_ domains are equally sensitive to DNMT1/DNMT3b loss. Interestingly, in the 5Aza-treated cells we find that all H3K9me3-HP1α/β domains show mild but uniform depletion of both Protect-seq signal, and HP1α and H3K9me3 levels (**Figure 5C** and **S.Figure 8**). To further highlight the local elimination (DKO) vs global depletion (5Aza-treated) of Protect-seq signal genome-wide, we color the E1-E2 projection of untreated HCT116 Hi-C data by Protect-seq signal in the three conditions (**Figure 5D**). From these data, we conclude that DNA methylation plays a complex role in maintaining heterochromatin.

Next, we ranked HCT116 B_2_ domains by H3K9me3 loss in DKO and split them into those that lose H3K9me3-HP1α/β status in DKO cells (disrupted domains) from those that retain it (persistent domains) (**Figure 5E** and **5F**). Based on our comparative analysis of B_2_ self-affinity (see above), we predicted that disrupted B_2_ domains will lose affinity for the remaining ones and fail to compartmentalize with them. To test this, we performed *in situ* Hi-C on HCT116, DKO, and 5Aza-treated cells. Indeed, in DKO cells we observe striking local defects in B_2_ compartmentalization (loss of checkering on the Hi-C map) and a global weakening of B_2_ compartmentalization in 5Aza-treated cells (**Figure 5G** and **5H**). These results reveal a strong connection between heterochromatin and 3D genome organization.

Next, we aimed to investigate the interaction profile acquired by disrupted domains in DKO. Aggregate analysis of contact frequency shows that disrupted domains change to a more neutral interaction profile (**Figure 5H**), reminiscent of the interaction profile of B_0_ domains. To address this in more detail, we examined the chromatin state at disrupted domains in DKO cells. We used available data to examine histone modifications H3K27me3, H3K9me2, H3K9me3, as well as H2A.Z in DKO cells (Lay et al., 2015; Spracklin & Pradhan, 2020). Indeed, persistent domains maintain a H3K9me3-HP1α/β chromatin state as expected. In contrast, disrupted domains transition into a chromatin state enriched for H3K9me2 and H2A.Z (**Figure 5I** and **S.Figure 8**), which is characteristic of B_0_ domains. In summary, these data suggest that disrupted domains in DKO cells display a similar Hi-C interaction profile and underlying chromatin state to B_0_ domains in HCT116.

Taken together, these results support our previous observations and suggest that H3K9me3 and HP1 are necessary for the spatial compartmentalization of constitutive heterochromatin in the nucleus. In addition, these data point to a DNA methylation-dependent plasticity in heterochromatin, where the chromatin states underlying B_0_ (poised) and B_2_ (constitutive) are transposable.

### The loss or gain of H3K9me3-HP1α/β heterochromatin does not alter replication timing

Our data suggests that upon loss of DNA methylation B_2_ domains can lose H3K9me3, HP1 and self-affinity and revert to a neutral inactive chromatin state. Replication timing has been proposed to maintain the global epigenetic state in human cells (Klein et al., 2021). In turn, histone deposition, HP1 proteins and DNMT1 are associated with chromatin restoration at the replication fork (Groth et al., 2007; Maison & Almouzni, 2004). Therefore, we hypothesized that the loss of late-replicating constitutive heterochromatin in DKO would be accompanied by a change in the timing of DNA replication at disrupted domains. To address whether replication timing is altered by the disruption of heterochromatin, we performed two-stage Repli-seq in HCT116 and DKO cells. Interestingly, we observe similar replication timing profiles between HCT116 and DKO cells (**Figure 6A**). Both cell lines have a similar number and size distribution of early and late replicating domains and these domains are highly overlapping (j=0.78), with minor differences (**S.Figure 9**).

**Figure 6:**
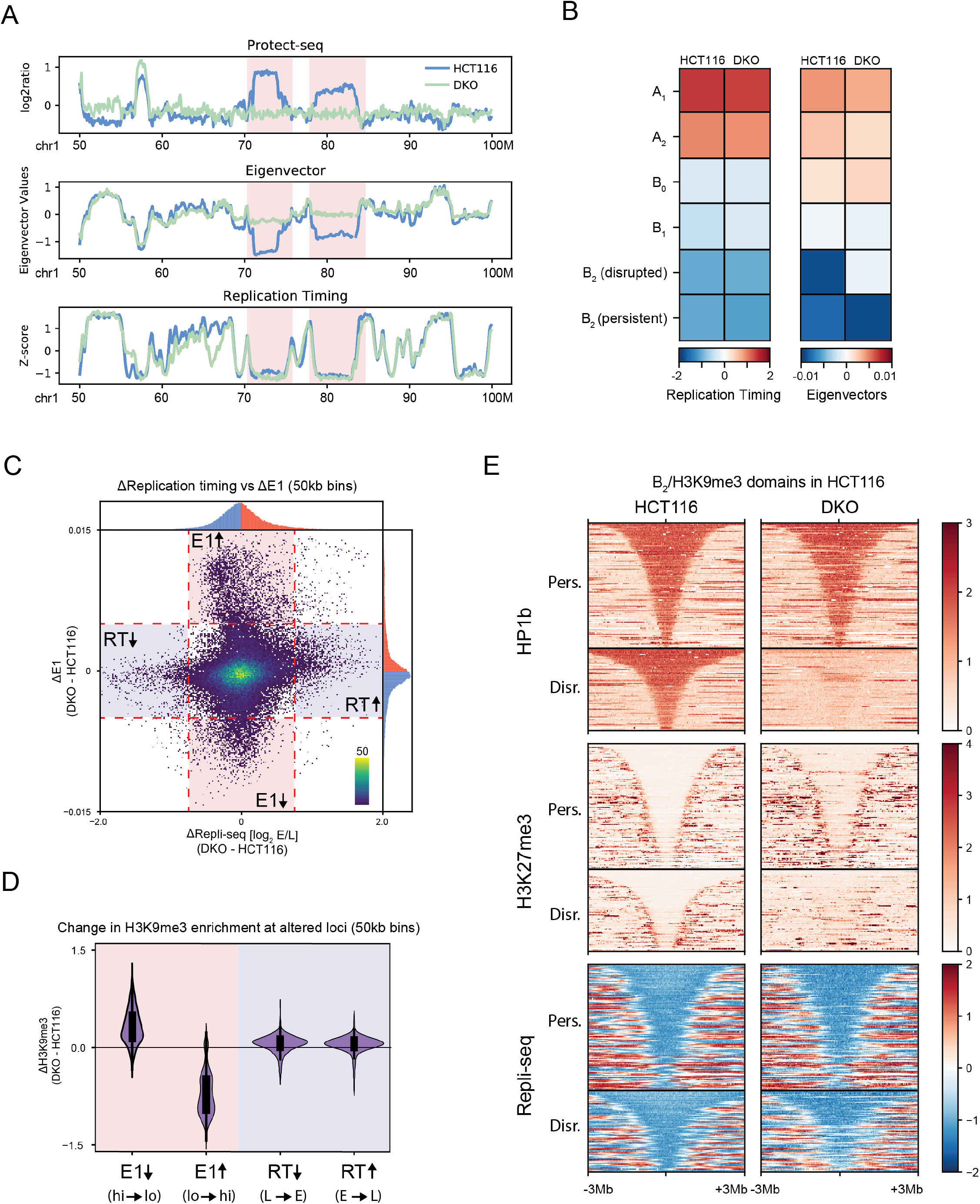
Loss or gain of H3K9me3-HP1α/β and replication timing alterations are uncorrelated. (A) Example region (chr1:50-100M) containing two disrupted domains in HCT116 cells (light blue) and DKO cells (light green) illustrating dramatic changes in compartmentalization without changes in replication timing. Top, Protect-seq signal track (log_2_ signal/input). Middle, eigenvector track (E1). Bottom, two-stage Repli-seq shown as z-score of log_2_(Early/Late). (B) Heatmaps of mean signal of Repli-seq (left) and E1 (right) over 50kb bins per interaction profile group in HCT116 and DKO. (C) Scatter plot of change in E1 score vs change in Repli-seq signal for 50kb bins (DKO - HCT116). Tail areas of uncorrelated variation of E1 and replication timing are gated and shaded. (D) Violin plots quantifying changes in H3K9me3 (DKO - HCT116) over groups of altered 50kb bins depicted in (C): decreased E1 score in DKO, increased E1 score in DKO, decreased Early/Late signal in DKO (delayed replication timing), increased Early/Late signal in DKO (hastened replication timing). (E) Stacked signal heatmaps of HP1β ChIP-seq, H3K27me3 ChIP-seq, and Repli-seq in HCT116 (left) and DKO (right) centered at persistent (top) and disrupted (bottom) B_2_ domains sorted vertically by size flanked by ±3Mb.

A fine scale analysis of individual loci further shows that changes in replication timing and changes in the Hi-C E1 eigenvector are uncoupled (**Figure 6B** and **6C**). Both persistent and disrupted B_2_ domains, which are late replicating in HCT116, remain late replicating in DKO cells (**Figure 6B** and **S.Figure 9**). Importantly, we do not see major replication timing differences within disrupted B_2_ regions (i.e. that lose H3K9me3 and HP1 and cease to compartmentalize in DKO cells) or within regions where H3K9me3 and HP1 was gained in DKO (**Figure 6A**, **6B, 6C, 6D**, and **S.Figure 9**). We further identified regions of differential replication timing and we find that those regions which show earlier replication timing in DKO correlate with loss in H3K27me3, but not H3K9me3. Together, our data suggest that the late timing of replication within B_2_ domains is not due to H3K9me3 and HP1 suppressing early onset of DNA replication.

In summary, we conclude that loss of DNA methylation has relatively minor effects on the population-averaged timing of DNA replication along the genome. Furthermore, replication timing is surprisingly insensitive to the presence or absence of H3K9me3-HP1α/β despite the role of the latter in B_2_ compartmentalization integrity.

### H3K9me3-HP1α/β heterochromatin suppresses CTCF binding sites

Our work thus far suggests that H3K9me3-HP1α/β domains co-segregate in the nucleus, permit loop extrusion but lack extrusion barriers. One striking observation in Hi-C data obtained with DKO and 5Aza-treated cells is the emergence of loop extrusion features in H3K9me3-HP1α/β domains, compared to HCT116 (**Figure 7A**). In other words, upon loss of DNA methylation, otherwise featureless B_2_ domains appear to acquire loop extrusion barriers. Moreover, we observe an increase in insulating loci in all interaction profile groups suggesting that this is not limited to H3K9me3-HP1α/β domains but rather is a global phenotype (**S.Figure 10**). Next, we aimed to understand the mechanism behind the gain of extrusion barriers.

**Figure 7:**
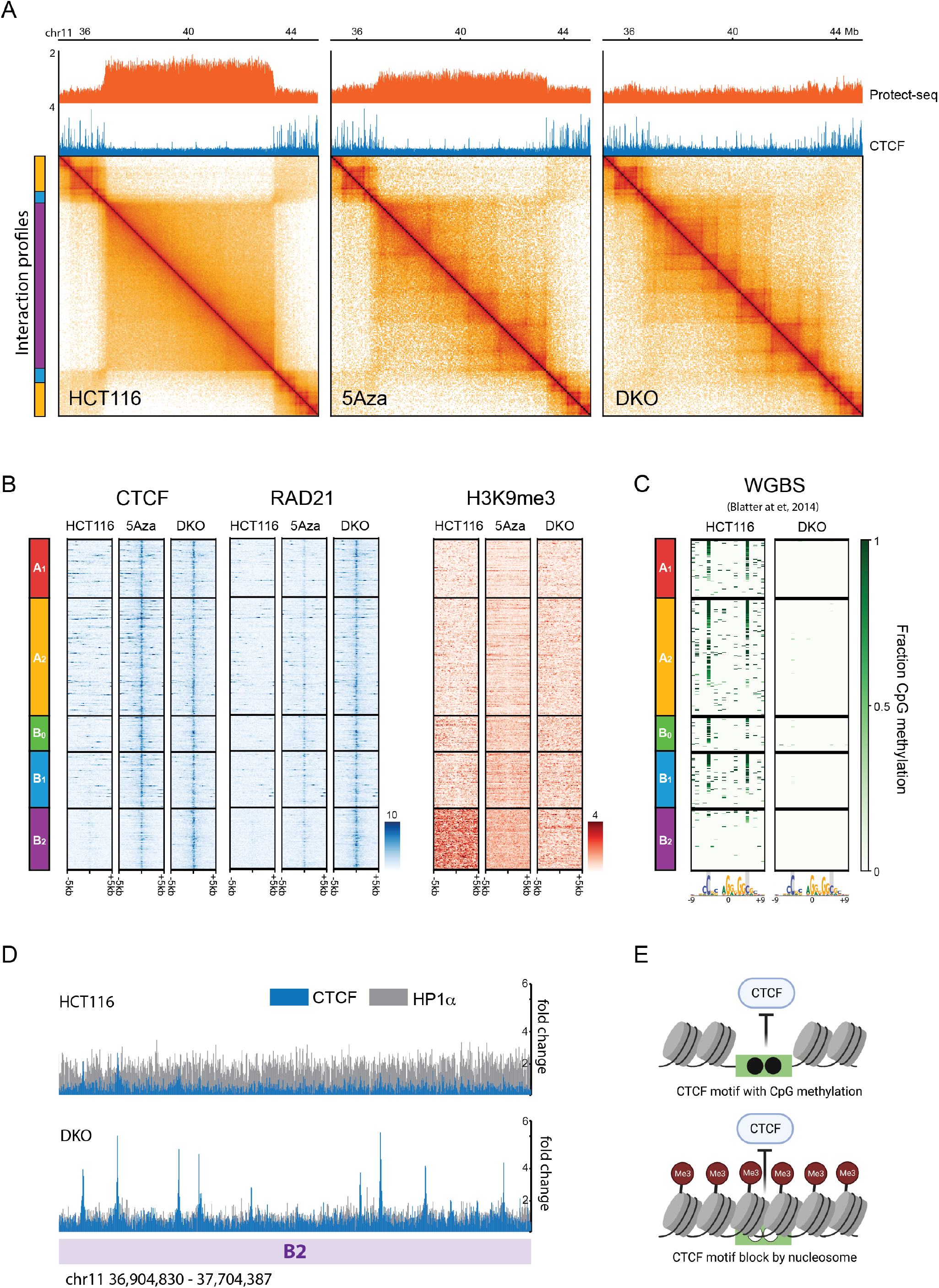
Two modes of CTCF binding suppression depend on DNA methylation. (A) Contact frequency maps of a 9-Mb genomic region (chr11:35.5-44.5M) in HCT116 (left), 5Aza (middle) and DKO (right) containing a representative example of reactivated CTCF sites. Top, Protect-seq and ChIP-seq track for CTCF. (B) Stacked heatmaps of reactivated CTCF sites for HCT116, 5Aza, and DKO cells centered on the CTCF motif displaying ChIP-seq signal for CTCF (left), Rad21 (middle), and H3K9me3 (right) flanked by ±5kb and segregated by interaction profile group. (C) Stacked heatmaps around reactivated CTCF site core motifs (19 bp) for HCT116, 5Aza, and DKO cells displaying fraction CpG methylation using whole genome bisulfite sequencing data from (Blattler et al., 2014). (D) Example of reactivated CTCF sites within a B_2_ domain (chr11:36.9-37.7M). Top, HCT116 ChIP-seq signal for CTCF (blue) and H3K9me3 (grey) overlayed. Bottom, DKO ChIP-seq signal for CTCF (blue) and H3K9me3 (grey) overlayed. (E) Model of two modes of CTCF regulation. Top, the direct mode involves CpG methylation within the core binding motif. Bottom, the indirect mode involves steric occlusion of the CTCF binding site by nucleosomes and/or other heterochromatic factors.

It has been shown that CTCF binding to DNA can be blocked by DNA methylation (Bell & Felsenfeld, 2000; Hark et al., 2000) and genome-wide loss of DNA methylation has been shown to increase CTCF occupancy at CpG-containing motifs (termed reactivated CTCF sites) (Maurano et al., 2015). Hence we hypothesized that new loop extrusion features seen in DKO and 5Aza-treated cells are due to reactivated CTCF sites. To confirm that loss of DNA methylation reactivates cryptic CTCF sites, we performed ChIP-seq in HCT116, DKO and 5Aza-treated cells. To identify high-confidence reactivated CTCF peaks, we chose overlapping reactivated CTCF peaks from DKO (this study), DKO (Maurano et al. 2015), and 5Aza (this study), not present in HCT116 (n=1050) (**S.Figure 10**). Reactivated CTCF sites are present in all interaction profile groups, consistent with our observation that the increase in extrusion barriers occurs globally (**Figure 7B**). In accordance with the role of CTCF as a barrier to loop extrusion, we also see an enrichment of cohesin complex factors RAD21 and SMC3 at reactivated CTCF sites only in DKO and 5Aza-treated cells (**Figure 7B** and **S.Figure 10**). To further demonstrate that reactivated CTCF sites are functional as extrusion barriers, we generated aggregate heatmaps of Hi-C contact frequency centered at reactivated CTCF sites for each interaction profile group (**S.Figure 10**). As expected, we observe an increase in insulation in DKO and 5Aza compared to HCT116. In sum, these data support that loss of DNA methylation leads to the emergence of functional CTCF sites which can act as barriers to stall loop extruding cohesin complexes.

To further investigate the genome-wide patterns of CTCF reactivation, we profiled DNA methylation, chromatin inaccessibility, and histone modifications in relation to our interaction profile groups. To our surprise, in normal untreated HCT116, CpG methylation at reactivated CTCF motifs is absent in B_2_ regions but present in all other groups (**Figure 7C** and **S.Figure 10**). These data suggest that DNA methylation could regulate CTCF via two mechanisms: direct and indirect. The direct mechanism relies on canonical CpG methylation within the core motif which is present in A_1_, A_2_, B_0_, B_1_ but not in the B_2_ interaction profile group. CTCF regulation via two methylated CpGs within the core motif has been extensively studied (Bell & Felsenfeld, 2000; Hark et al., 2000; Hashimoto et al., 2017; Maurano et al., 2015; Renda et al., 2007; Wang et al., 2012). The indirect mode of regulation is likely independent of CpG methylation within the motif and is widespread in B_2_. Consistent with this observation, CTCF motifs within B_2_ contain fewer CpG dinucleotides than the canonical core motif (**S.Figure 10**). We speculate that this mechanism acts through nucleosome occlusion, which is consistent with the strong histone H3 and HP1α/HP1β signal directly over the CTCF motif (**Figure 7D** and **S.Figure 10**). In agreement with our results, a similar 5mC/nucleosome occlusion model has been proposed to regulate CTCF in mouse embryonic stem cells (Teif et al., 2014; Wiehle et al., 2019)

In summary, our data reveals two mechanisms by which CTCF occupancy may be modulated in the genome: (i) via CpG methylation of the CTCF core motif, likely to block zinc-finger binding and (ii) via H3K9me3-HP1α/β heterochromatin, which occludes the binding of CTCF through a mechanism that does not involve CpG methylation of CTCF motifs, but is dependent on DNA methyltransferases and/or repressive histone modifications in the region (**Figure 7E**). We further demonstrate that reactivated CTCF binding in B_2_ is functional (i.e. act as loop extrusion barriers), providing additional support to our previous observation that B_2_ regions permit active loop extrusion.

## Discussion

We performed a detailed investigation of nuclear compartmentalization and its relationship to chromatin state. In HCT116 cells, this led to the discovery of multiple distinct inactive compartmental domain types. We applied an unsupervised dimensionality reduction approach that uses interchromosomal contacts to identify sets of loci with comparable interaction patterns. No discrete segmentation can likely fully capture the variability in long-range contact frequency profiles observed in even very deeply sequenced contact frequency maps. However, by integrative comparison with other genomic and functional data, we were able to disentangle the influence of genomic position and find that interaction profiles can partition loci into at least five groups that, by and large, share common chromatin state features. The A_1_ and A_2_ groups represent transcriptionally active loci with A_1_ likely representing nuclear speckle-associated domains. The three remaining interaction profile groups are attributable to transcriptionally inactive chromatin states and low gene density. One group, B_0_, appears to be in a poised state and is enriched in H3K9me2 and H2A.Z. Another group, B_1_, exhibits H3K27me3/EZH2 enrichment and intermediate DNA methylation. Finally, group B_2_ is defined by trimethylation of H3K9 simultaneously co-occupied by factors HP1α and HP1β and spans long, continuous stretches of hypomethylated blocks of DNA (also known as partially-methylated domains) of up to several megabases. Our results show that each inactive interaction profile group exhibits a distinct chromatin state, Protect-seq signal, DNA methylation status, and display differences in homotypic affinity and the regulation of loop extrusion barriers.

The H3K9me3-HP1 chromatin state shows a strong self-interaction tendency that correlates with signal intensity and shows a wide diversity in genomic distribution across many cell types. A similar self-interaction tendency is observed for A_1_ loci in *trans*. This suggests that in the case of B_2_, and A_1_ to some extent, the interaction profile does indeed reflect strong spatial clustering and compartmentalization of loci. Such spatial segregation can be explained by polymer theory. Attraction based on even a subset of chromatin marks or associated factors can drive compartmentalization, where differential biomolecular affinities lead to microphase separation (de Gennes, 1979; Leibler, 1980; Matsen & Schick, 1994), one potential embodiment being the formation of condensates (Erdel & Rippe, 2018; Shin & Brangwynne, 2017). In block copolymer systems, compartmentalization as an emergent property will depend on the (i) abundance or volume fraction of monomer types (e.g. A/B ratio) and (ii) the lengths and distribution of homotypic blocks (e.g. AAAABBBB vs ABABABAB) and (iii) the nature of their multivalent biomolecular interactions and incompatibilities (Erdel & Rippe, 2018; Gavrilov et al., 2011; Halperin, 1991; Singh et al., 2020). Importantly, compartmentalization patterns in population Hi-C maps will not only be influenced by these intrinsic biomolecular properties but also by cell-cell or allelic heterogeneity in chromatin state, and potentially by tethering of different regions to the lamina and nuclear bodies. Our results on B_2_ are in close agreement with a recent study of the relationship between chromatin marks and Hi-C, which considers an attraction-repulsion model of compartmentalization and suggests that GM12878, with mostly narrow H3K9me3 domains, and HCT116, displaying large and abundant H3K9me3 domains, likely lie on two ends of a spectrum of H3K9me3 distribution (Nichols & Corces, 2021). In some cell types, the H3K9me3-HP1 chromatin state is limited to shorter, more dispersed regions, contributing little to global compartmental segregation. Nevertheless, many cell types, including H1 and GM12878, have H3K9me3-HP1α associated chromatin at domains coating ZNF gene clusters on chromosome 19 (Vogel et al., 2006) that do exhibit obvious preferential homotypic interactions. In cell types with prevalent heterochromatin, we propose that H3K9me3-HP1α/β chromatin exerts a dominant organizing force (e.g. in HCT116 and inverted Rod photoreceptor nuclei (Falk et al., 2019; Solovei et al., 2009)), by forming spatially segregated clusters of chromatin domains.

Our work demonstrates a remarkable intracellular and cell-cell diversity in inactive chromatin and its relationship to 3D genome organization. The existence of cell-type specific chromatin and contact frequency profiles highlights the need to apply unsupervised methods for *de novo* assessment of any given cell type. Surprisingly, our approach identified an interaction profile group, B_0_, not found in GM12878 cells. In HCT116 cells, B_0_ forms large domains that display a neutral interaction preference, meaning these domains do not display strong homotypic interactions. The B_0_ interaction profile group overlaps the ‘D’ cluster reported on chromosome 4 in (Nichols & Corces, 2021) in HCT116 cells. Yet another type of inactive chromatin appears to underlie the majority of loci labeled B2/B3 in GM12878 and remains poorly characterized. In GM12878 lymphoblastoid cells, the primary features of B2/B3 that were reported to be enriched were Nucleolus-associated domains (NADs) and Lamina-associated domains (LADs); however, these assays were performed in dissimilar cell types: HeLa (Németh et al., 2010), HT1080 fibrosarcoma (van Koningsbruggen et al., 2010) and skin fibroblasts (McCord et al., 2013). Detecting novel interaction profiles in other cell types and elucidating the molecular intermediates underlying affinity-mediated contact patterns will require a combination of unsupervised techniques and deep chromatin profiling (Boyle et al., 2020; Rosencrance et al., 2020; Zhang et al., 2020).

Our results reveal striking connections between DNA methylation, H3K9me3 and HP1 deposition, and 3D chromosome organization at the level of chromosome compartmentalization and loop extrusion. DKO cells (possessing a deletion of DNMT3B and a DNMT1 hypomorph (Rhee et al., 2002)) have H3K9me3-HP1 defects (Lay et al., 2015; Spracklin & Pradhan, 2020) and acute inhibition of DNA methyltransferases leads to global heterochromatin depletion. We demonstrate that the heterochromatic state is integral to its nuclear compartmentalization. Interestingly, when DNA methylation is lost, H3K9me3-HP1/B_2_ domains convert to the B_0_ inactive chromatin state that lacks self-affinity yet maintains late replication timing. These data lead us to speculate that B_0_ domains represent a poised inactive chromatin state. These domains also acquire H2A.Z and we propose that the presence of this histone variant could function to interfere with heterochromatin deposition and spreading (e.g. the higher-order structure of the heterochromatic nucleosomal fiber may be incompatible with H2A.Z), as has been shown in budding yeast (Meneghini et al., 2003). Moreover, these results point to an epigenetic plasticity governing dramatic changes in nuclear organization that may be essential for differentiation, lineage commitment and cell decision making, and may become dysregulated in cancer and aging.

Loop extrusion and compartmentalization shape different aspects of genome organization. While the forces driving compartmentalization are believed to be global and intimately linked to the state of chromatin, the degree to which loop-extruding cohesins are influenced by the epigenome is not well understood. Here we find that H3K9me3-HP1α/β heterochromatin, exemplified by the B_2_ interaction profile group in HCT116, is permissive to loop extrusion by cohesin yet displays a striking absence of loop extrusion features (stripes and dots) because it intrinsically resists the binding of CTCF. As loop extrusion has been shown to reduce the strength of compartmentalization and interfere with the segregation of short compartmental domains (Nuebler et al., 2018; Rao et al., 2017; Schwarzer et al., 2017; Wutz et al., 2017), our results represent a complementary phenomenon: strongly compartmentalizing heterochromatin suppressing the imposition of extrusion barriers (CTCF-bound sites). These results highlight the two-way interplay between compartmentalization and extrusion.

Loss of DNA methylation, or DNA methyltransferases, allows dormant CTCF sites to become bound by CTCF and act as loop extrusion barriers. This occurs in at least two ways depending on chromatin domain type. The predominant mechanism for CTCF binding suppression, except in B_2_, is DNA methylation within the core CTCF motif. Interestingly, in B_2_ domains CTCF binding is also prevented in a DNA methylation-dependent manner but not through CpG methylation of the motifs themselves. We propose that in B_2_ domains DNA methylation at other sites, possibly near the binding sites, contributes to nucleosome positioning over CTCF motifs which in turn block CTCF binding. Loss of DNA methylation then allows opening chromatin locally, contributing to the B_2_-to-B_0_ transition and CTCF binding.

Why is heterochromatin so diverse, both within and between cell types? The classic definition of heterochromatin originated from staining mitotic chromosomes (Heitz, 1928) and later came to be associated with histone modifications (Trojer & Reinberg, 2007). We now have a more nuanced understanding of the molecular details including several types of repressive histone modifications and associated proteins and their genomic distributions across cell types. However, the function of combinatorial modifications, molecular interactions between heterochromatin factors, and their contribution to genome organization remains less clear. Our work begins to unravel the diversity and plasticity in inactive chromatin and its influence on genome compartmentalization, nuclear architecture, and other chromosome organizing processes (loop extrusion, replication, DNA repair, cell division) to accomplish biological functions.

## Acknowledgements

We thank Pierre-Olivier Estève, Anton Goloborodko, Gretel Edgeworth, E. M. Breville, and members of the Dekker and Mirny labs for helpful insights and discussion. We thank Kirill Polovnikov for advice on spectral clustering. We thank Nicki Fox, Johan Gibcus, and Geoffrey Fudenberg for critical reading of the manuscript. This work was supported by New Englands Biolabs, Inc. and a grant from the National Institutes of Health Common Fund 4D Nucleome Program to J.D. and L.A.M (U54-DK107980 and UM1-HG011536). J.D. is an investigator of the Howard Hughes Medical Institute.

## Author Contributions

G.S. and S.P. conceived the study. G.S. designed and performed experiments. N.A., G.S., and M.I. performed data analysis. M.I. and N.C. performed polymer simulations. All authors contributed to data interpretation. G.S., N.A., L.A.M., and J.D. wrote the manuscript.

## Data Availability

The references and accession numbers of published data used and analysed in this work are indicated in **Supplementary Table** 2. All data sets generated in this study are deposited in the NCBI Gene Expression Omnibus (GEO; http://www.ncbi.nlm.nih.gov/geo/) under the accession number GSEXXXXXX.

---

**S.Figure 1:**
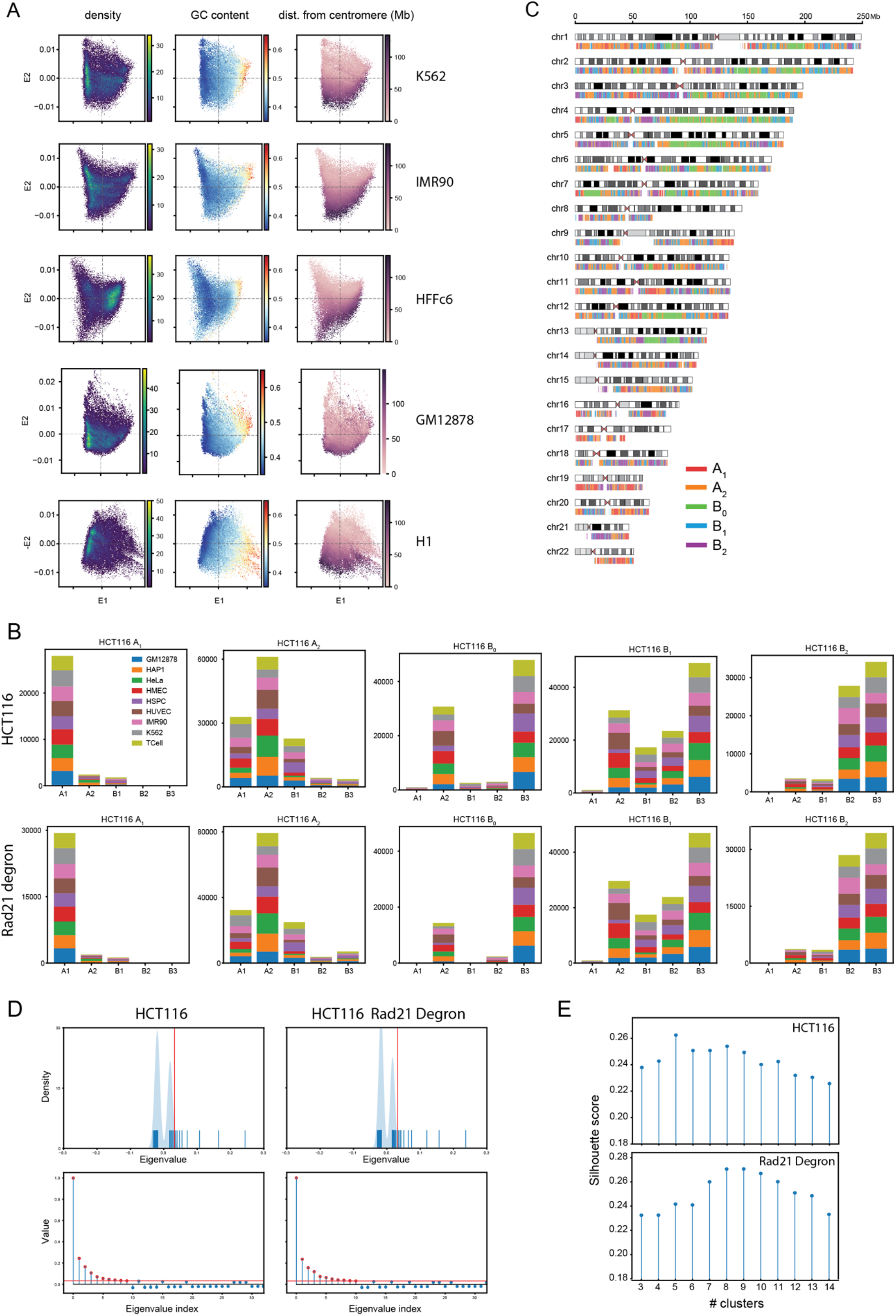
Spectral decomposition and clustering in HCT116. (A) E1 vs. E2 scatter plots of 50kb genomic bins from five additional cell types (K562, IMR-90, HFFc6, GM12878, H1-hESC) colored by point density (left), GC content (middle), and distance from centromere (right). (B) Distributions of SNIPER subcompartment labels assigned to genomic bins in each interaction profile group across nine other cell types for HCT116 (top) and HCT116 RAD21-degron (bottom). (C) Ideogram plot of interaction profile groups in HCT116 (A_1_, red; A_2_, yellow; B_0_, green; B_1_, blue; B_2_, purple). (D) Top, rug plot of the leading 128 eigenvalues for HCT116 (left) and HCT116 RAD21-degron (right). Vertical red line indicates the eigenvalue cutoff. Bottom, same eigenvalues plotted in descending order of absolute value. Eigenvalues corresponding to retained vectors used for clustering are indicated in red. (E) Silhouette scores calculated for *k*-means clustering on eigenvectors from HCT116 (top) and HCT116 RAD21-degron (bottom) as a function of the number of clusters, *k*.

**S.Figure 2:**
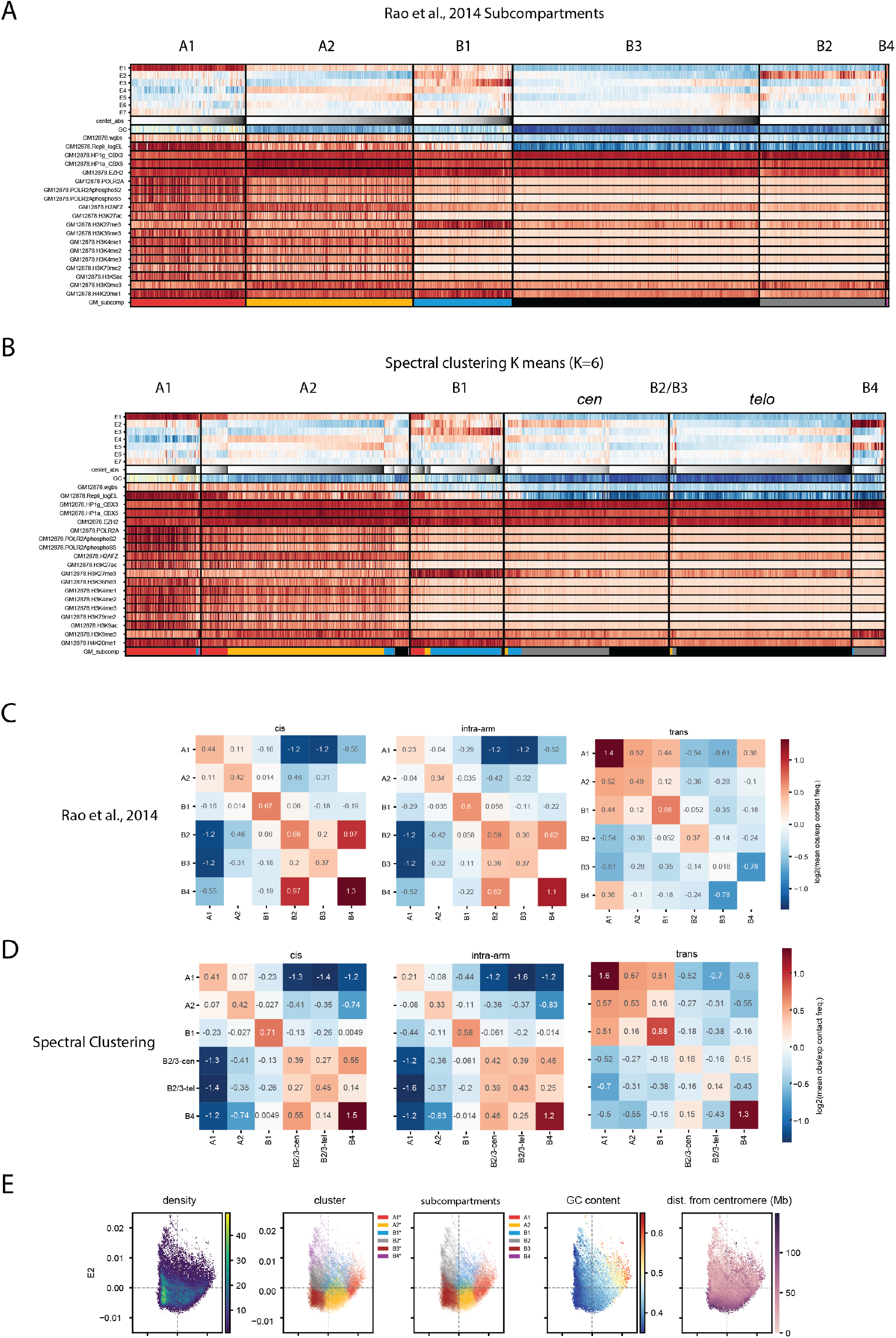
Spectral decomposition and clustering in GM12878. (A) Feature heatmap for GM12878 based on 6-subcompartment labels from (Rao et al., 2014). The tracks displayed are the seven leading eigenvectors (E1-E7), GC content, fraction CpG methylation, replication timing (Early/Late), and ChIP-seq for a range of factors and histone modifications. Columns (50-kb bins) within each subcompartment are sorted by distance from centromere. Colors are assigned to the subcompartment labels in the last row (A1: red, A2: yellow, B1: blue, B2: grey, B3: black). (B) Feature heatmap for GM12878 based on spectral clustering of E1-E7 (*k*=6). Rows display the same tracks as in (A). Columns within each cluster are sorted first by subcompartment label assignment, then by distance from centromere. The last row assigns a color to each bin based on its subcompartment label as in (A) (A1: red, A2: yellow, B1: blue, B2: grey, B3: black). Names are assigned to the clusters based on similarity to (A). The main differences with Rao et al., 2014 subcompartment assignments are (1) a more balanced division between B2 and B3 based on centromere/telomere proximity and (2) an expanded sixth cluster, B4*, that acquires B3 loci having highly enriched H3K9me3 and HP1γ. (C) Heatmaps of pairwise mean observed/expected contact frequency between subcompartments in (Rao et al., 2014) based on *cis* (left), intra-arm (middle), and *trans* (right) contacts. (D) Heatmaps of pairwise mean observed/expected contact frequency, as in (C), but between spectral clusters from (B). (E) E1 vs. E2 scatter plots from GM12878 colored by point density, GC content, spectral cluster label, subcompartment label, and distance from centromere.

**S.Figure 3:**
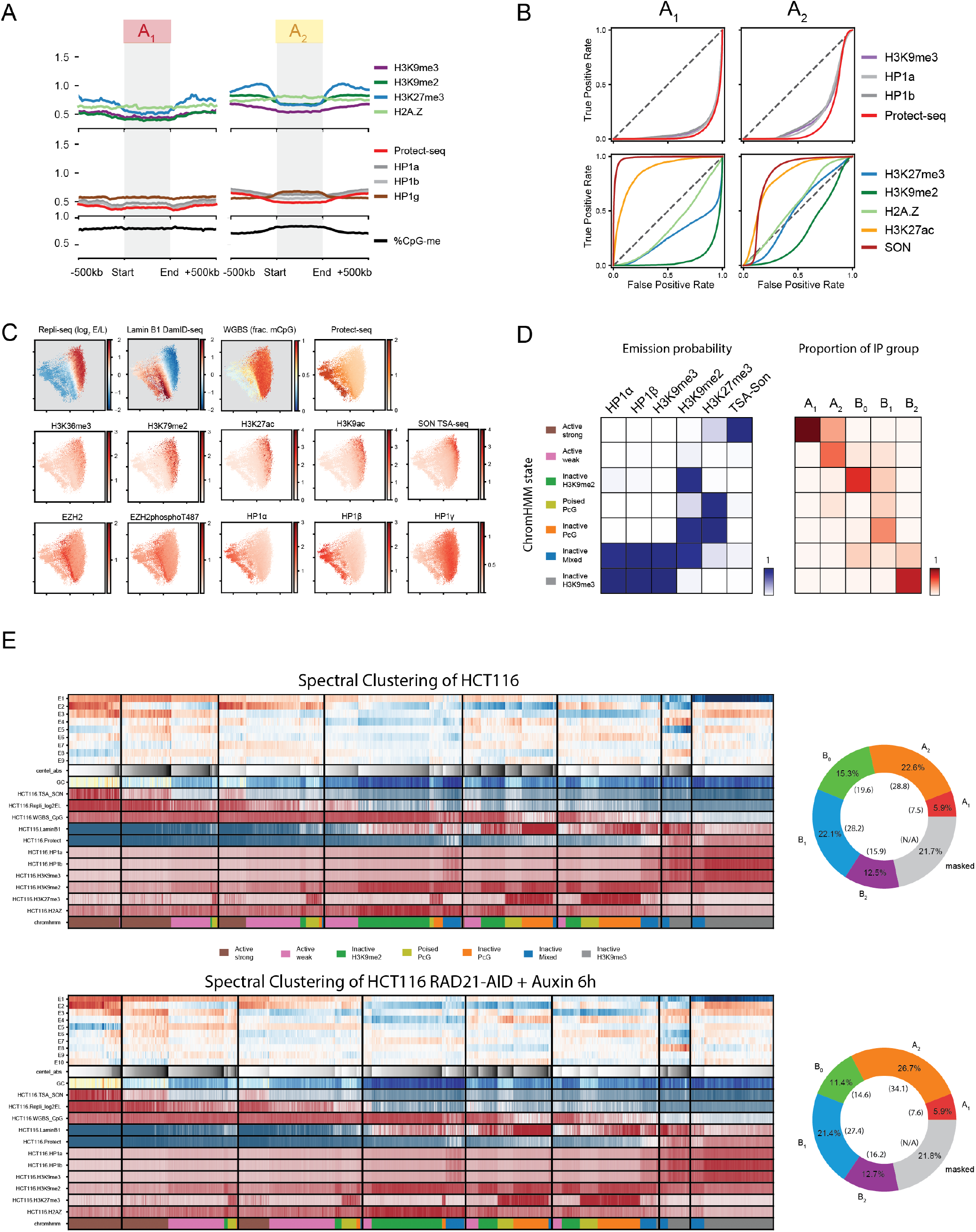
Chromatin state composition of interaction profile groups in HCT116. (A) Metaplots displaying signal enrichment for the same features as Figure 2C for A_1_ and A_2_ domains. (B) ROC curves assessing the prediction performance of individual 50kb-aggregated functional tracks as binary classifiers as in Figure 2D but for A_1_ and A_2_ loci. Additionally, curves for active marks (ChIP-seq for H3K27ac and TSA-seq for SON) are shown. (C) E1 vs. E2 scatter plots of 50-kb bins colored by point density and ChIP-seq for various factors and histone modifications. (D) Left, emission probabilities for ChromHMM model on five ChIP-seq for repressive marks and SON (TSA-seq for nuclear speckle marker) trained on 50kb bins. Right, heatmap showing the distributions of ChromHMM state labels found in each interaction profile group (columns). (E) Left, feature heatmaps for spectral clustering on HCT116 (top) and the cohesin-depleted HCT116 RAD21-AID line from (Rao et al., 2017) (bottom). The tracks displayed are the same as in Figure 1D but also include various histone marks. Columns (50-kb bins) within each cluster are sorted first by ChromHMM state (as per the model in (D)) and then by distance from centromere. The last row assigns a color to each bin based on its ChromHMM state. Right, donut plots showing hg38 percentage covered by each interaction profile group (top, HCT116; bottom, HCT116 RAD21-AID). Note: translocations and unmappable areas are masked. Percentages excluding translocations and unmappable areas are in parentheses.

**S.Figure 4:**
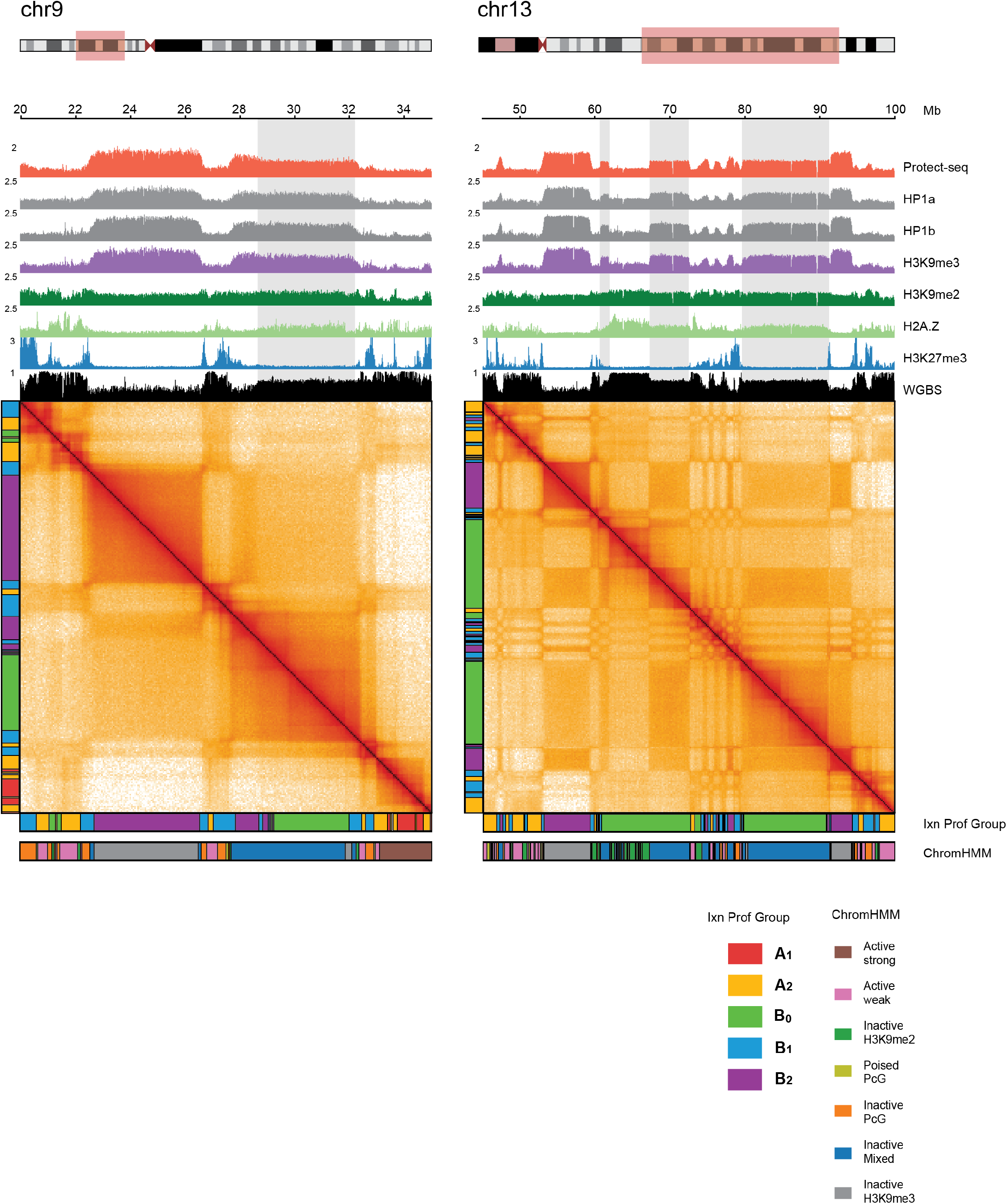
Examples of mixed-state domains (constitutive/poised). Two example regions that display a mixed chromatin state (chromHMM) and Hi-C signal (B_0_ and B_2_). Highlighted boxes illustrate domains with fractional heights relative to neighboring domains (Protect-seq, ChIP-seq, WGBS). Note the faint appearance of loop extrusion features in the Hi-C maps as well.

**S.Figure 5:**
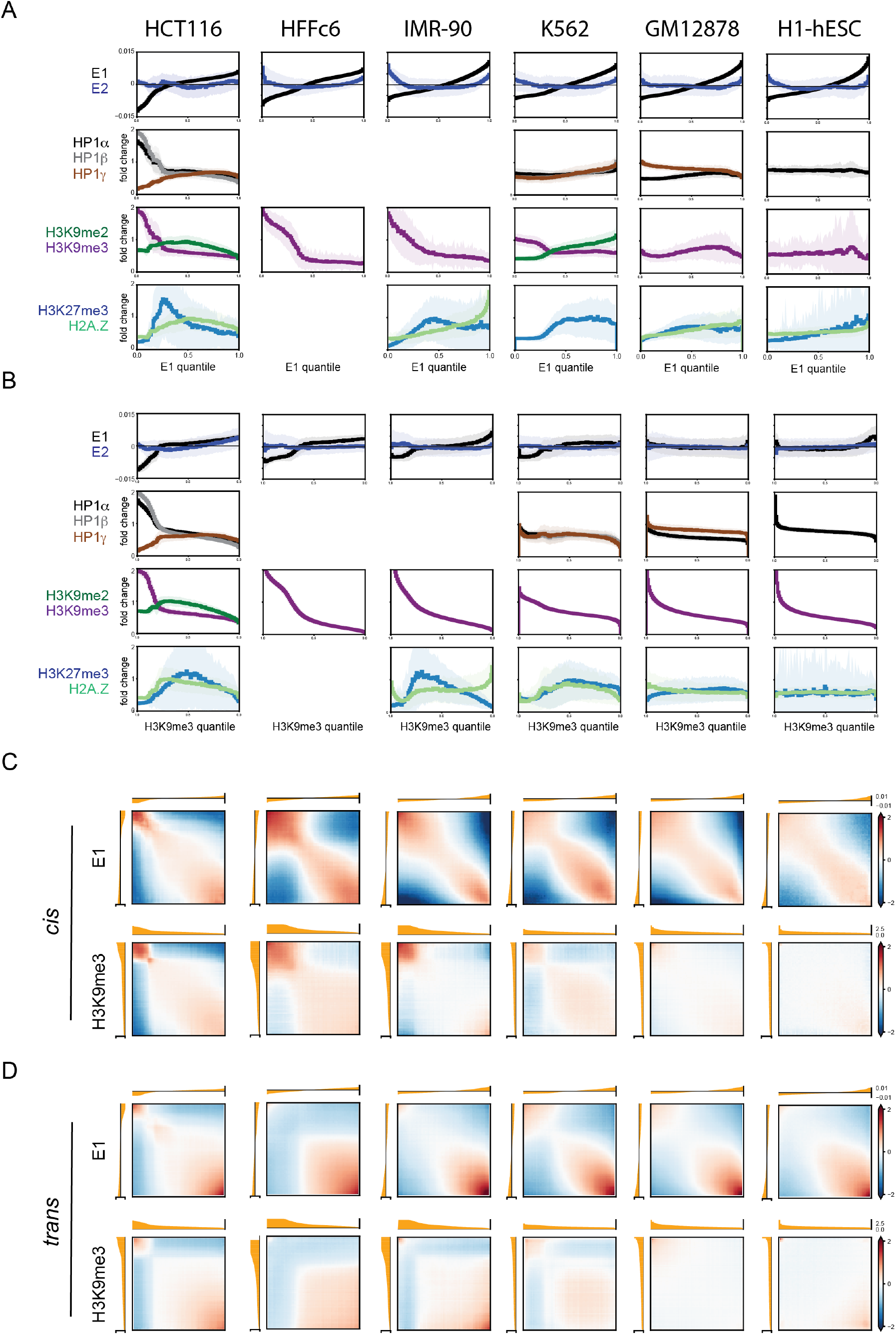
Comparative analysis of compartmentalization and heterochromatin marks. Comparative analysis of genome organization and heterochromatic marks across HCT116, HFFc6, IMR90, K562, GM12878 and H1-hESC. (A) Histograms of ChIP-seq signal for repressive histone marks as in Figure 3A based on eigenvector (E1) percentile and displayed in ascending order of E1 rank. Includes additional histograms for E1 and E2 (top) and data for two additional cell types: lung fibroblasts IMR-90 and foreskin fibroblasts HFFc6. (B) Histograms of ChIP-seq signal for repressive histone marks as in Figure 3D based on H3K9me3 percentile and displayed in descending order of H3K9me3 rank. Includes additional histograms for E1 and E2 (top) and data for IMR-90 and HFFc6. (C) Bivariate histograms of *cis* observed / expected contact frequency as in Figure 3B and 3C based on E1 percentile in ascending order (top) and H3K9me3 percentile in descending order (bottom). (D) Bivariate histograms as in (C) but describing observed / expected contact frequency in *trans*.

**S.Figure 6:**
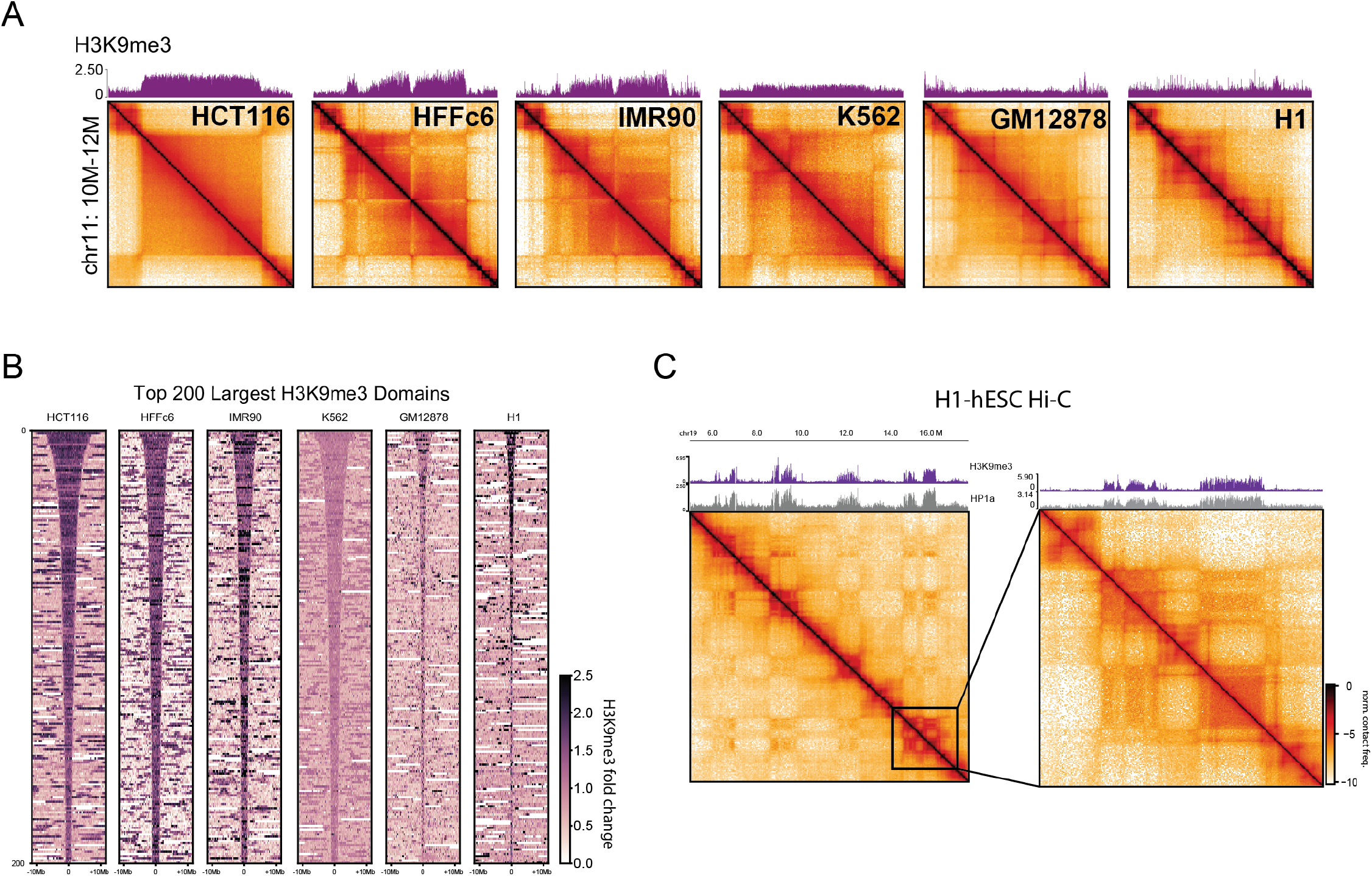
Comparative analysis of H3K9me3 domains. Comparative analysis of genome organization and heterochromatic marks across HCT116, HFFc6, IMR90, K562, GM12878 and H1-hESC. (A) Expanded example domain across cell types as in Figure 3E including data for IMR-90 and HFFc6. (B) Stacked signal heatmaps centered at the top 200 largest H3K9me3 domains detected in six cell types displaying H3K9me3 signal. (C) Example of homotypic interactions at H3K9me3-HP1α domains on chr19 in H1-hESC.

**S.Figure 7.**
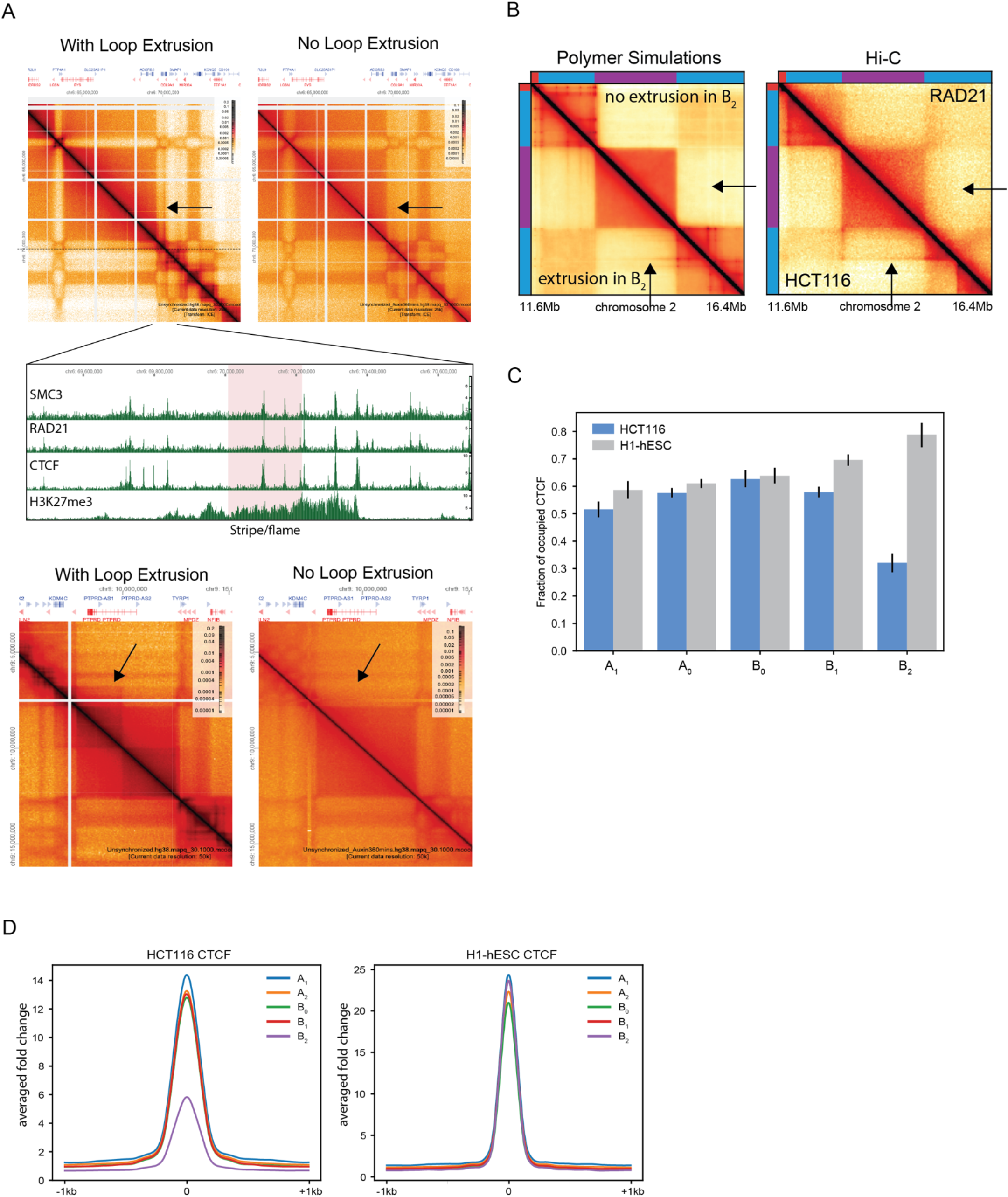
Evidence of loop extrusion but lack of CTCF within heterochromatin domains. (A) Two examples of cohesin-dependent loop extrusion features traversing a B_2_ domain. Hi-C maps of HCT116 (left columns) and HCT116-RAD21 auxin depletion (right columns). Arrows indicate loop extrusion features that are dependent on the cohesin complex: stripe (top Hi-C map) and TAD (bottom Hi-C map). Middle panel, ChIP-seq tracks of SMC3, RAD21, CTCF, and H3K27me3 for the stripe (highlighted in pink) and surrounding region (B) Contact frequency maps from *in silico* polymer simulations (left) compared to experimental Hi-C (right). Arrows indicate a stripe next to a B_2_ domain that extends parallel to its edge in HCT116. Experimental data is replicated when cohesin traversal is permitted (lower triangle) and does not appear when loop extrusion is blocked at the B_2_ domain (upper triangle). (C) Fraction peaks detected at all known CTCF sites (from Maurano et al., 2015) occupied in HCT116 and H1-hESC ChIP-seq grouped by HCT116 interaction profiles. (D) Average fold enrichment of CTCF ChIP-seq across all known CTCF sites used in (C) for HCT116 and H1-hESC.

**S.Figure 8:**
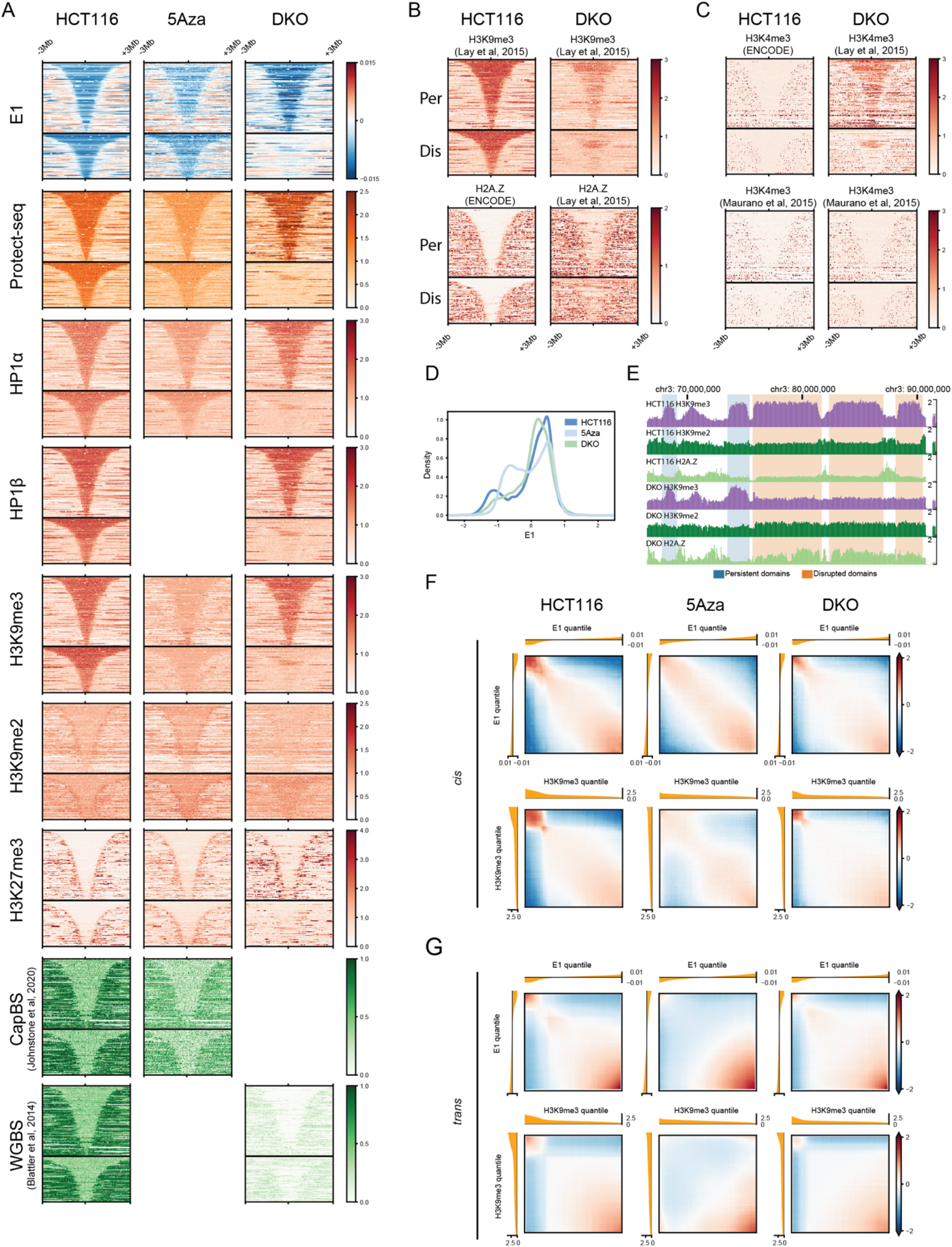
Maintenance of H3K9me3-HP1α/β heterochromatin depends on DNA methylation homeostasis. (A) Stacked signal heatmaps centered at persistent and disrupted B_2_ domains (not scaled) displaying various signal tracks in HCT116, 5Aza-treated cells, and DKO. Hybrid Selection Capture BS DNA methylation (CapBS) data were obtained from (Johnstone et al., 2020) and WGBS data were obtained from (Blattler et al., 2014). (B) Stacked signal heatmaps centered at persistent and disrupted B_2_ domains identified in this study displaying H3K9me3 and H2A.Z signal from (Lay et al., 2015). Note that the H3K9me3 domains in the DKO line used in that study appear slightly divergent from those detected here. (C) Stacked signal heatmaps similar to (B) but displaying H3K4me3 ChIP-seq from (Lay et al., 2015) and (Maurano et al., 2015). The first study shows a remarkable DKO-specific co-enrichment of H3K4me3 signal with H3K9me3 marking persistent domains, but this result was not reproduced in (Maurano et al., 2015). (D) KDE plots of E1 signal in HCT116, 5Aza-treated cells, and DKO. (E) Example region (chr3:70-90M) showing persistent (blue shading) and disrupted (orange shading) domains. ChIP-seq tracks for H3K9me2, H3K9me3, and H2A.Z in HCT116 (top 3 tracks) and DKO (bottom 3 tracks) (F) Bivariate histograms of *cis* observed / expected contact frequency based on E1 percentile (top) and H3K9me3 percentile (bottom) in HCT116, 5Aza-treated cells, and DKO. (G) Same as (F) but for *trans* contact frequency in HCT116, 5Aza-treated cells, and DKO.

**S.Figure 9:**
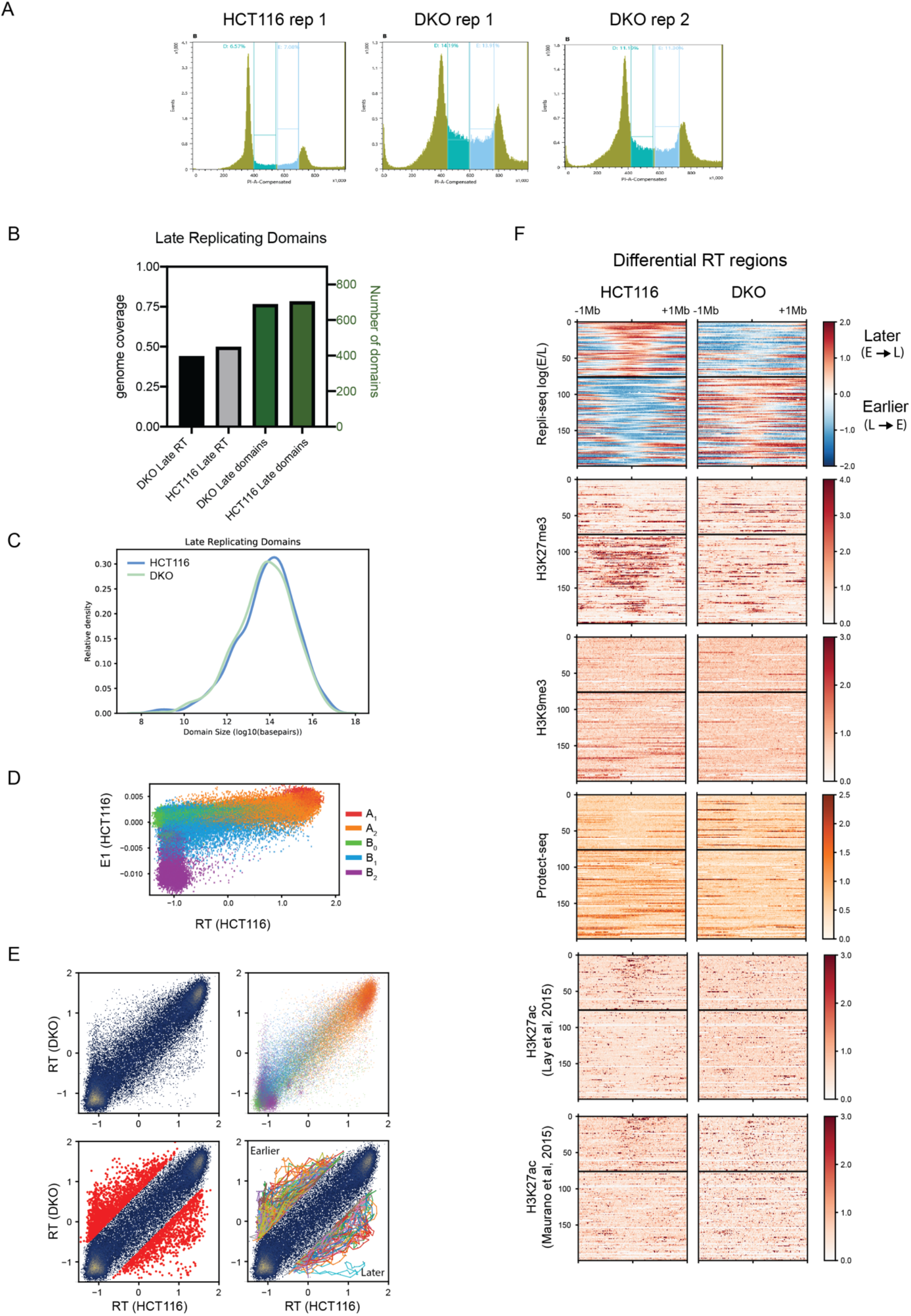
Altered replication timing does not occur at loci that lose or gain H3K9me3-HP1α/β but is enriched at loci that lose H3K27me3. (A) HCT116 (left), 5Aza-treated (middle), and DKO (right) cells were stained with propidium iodide and FACS sorted (SONY SH-800) based on DNA content (early S v late S). (B) Total number (green) and genome coverage (black) of late replicating domains detected in HCT116 and DKO using a Gaussian HMM. (C) KDE plots of domain size of late replicating domains (log10) in HCT116 and DKO. (D) Scatter plot of eigenvector (E1) and replication timing (RT) signal in HCT116 at 50-kb resolution colored by interaction profile label (A_1_ (red), A_2_ (orange), B_0_ (green), B_1_ (violet), B_2_ (blue)). (E) Differential replication timing analysis. Top: Left, scatter plot of 50-kb genomic bins based on z-scored Repli-seq log2(Early/Late) in HCT116 vs DKO. Right, same scatter plot colored by interaction profile label. Bottom: Left, same scatter plot with loci exhibiting a change >= 0.75 highlighted in red. Right, same scatter plot with continuous merged differential regions connected using colored lines. (F) Stacked signal heatmaps centered at differentially replicating regions (not scaled) divided into later/delayed onset (top) and earlier/hastened onset (bottom) regions displaying various signal tracks in HCT116 and DKO cells (n=199).

**S.Figure 10:**
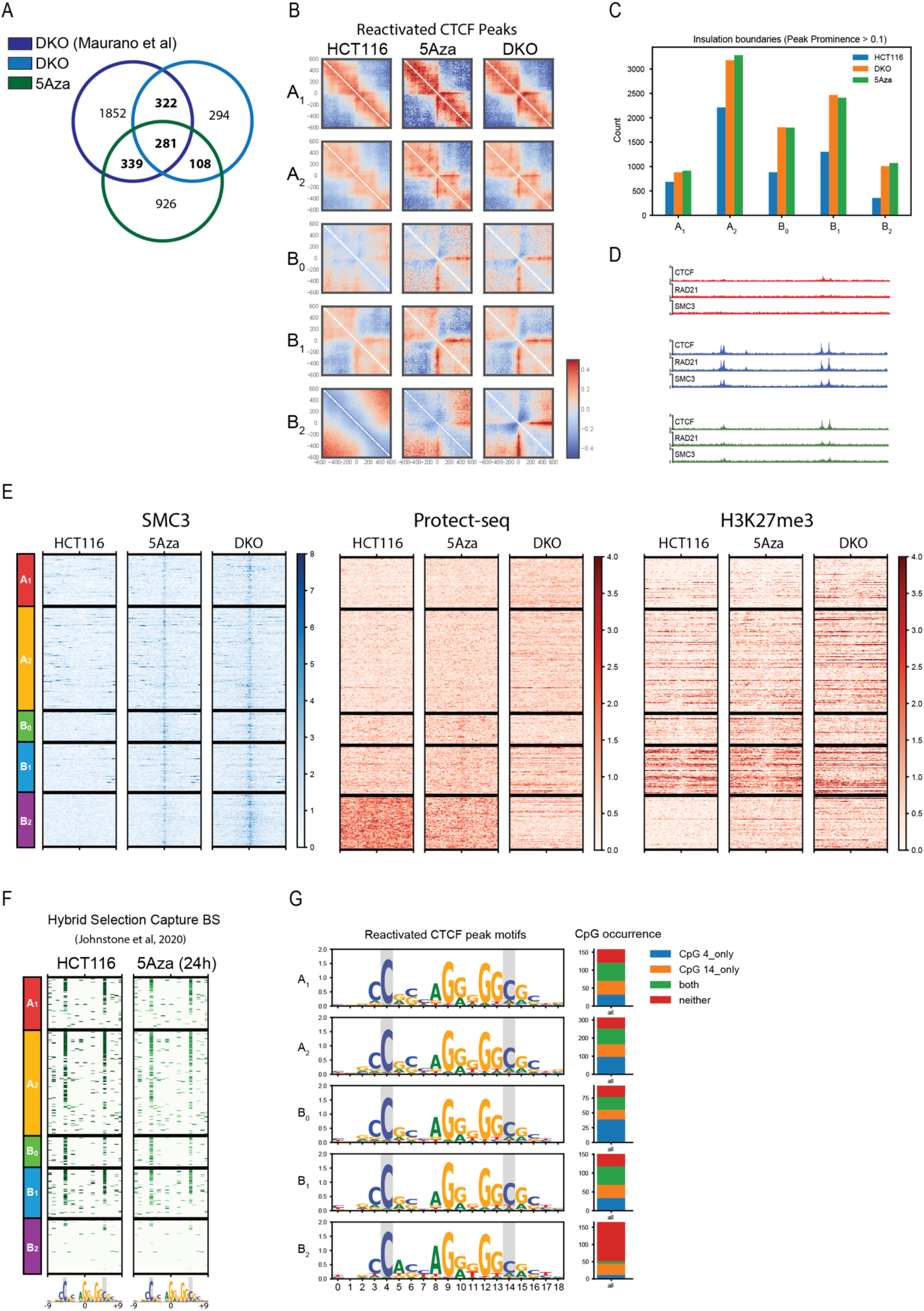
Reactivated CTCF sites. (A) Venn diagram of CTCF peaks in DKO (this study), 5Aza (this study), and DKO (Maurano et al., 2015). Union between CTCF peaks used to define reactivated CTCF sites. (B) Average observed/expected Hi-C maps around reactivated CTCF binding sites within each interaction profile group, centered at CTCF motifs oriented as indicated in HCT116 (left), 5Aza (center), and DKO (right) cells. (C) Quantification of total number of insulating loci with peak prominence score > 0.1 per interaction profile group. (D) Example region (chr11:39-40M) of reactivated CTCF sites blocking cohesin (RAD21 and SMC3). (E) Stacked heatmaps of reactivated CTCF sites for HCT116, 5Aza, and DKO cells centered on the CTCF motif displaying ChIP-seq signal for SMC3 (left), Protect-seq (middle), and H3K27me3 (right) flanked by ±5kb and segregated by interaction profile group. (F) Similar to Figure 7C. Stacked heatmaps around reactivated CTCF site core motifs (19 bp) for HCT116 and 5Aza-treated cells displaying fraction CpG methylation using hybrid selection capture bisulfite sequencing data from (Johnstone et al., 2020). (G) Left: sequence logos for the reactivated CTCF motifs in each interaction profile group. Right: frequencies of CpG occurrence at motif positions 4 and 14 in each set of reactivated CTCF sites. Note: nucleotides 4 and 14 depend on the motif start, other publications refer to these CpG nucleotides as 2 and 12 (e.g. Hashimoto et al., 2017) or 1 and 11 (e.g. Wang et al., 2012).

## Methods

### Cell culture

HCT116 and DKO cells were cultured in McCoy5A media. DKO cells were grown in the presence of G418, geneticin. All media was supplemented with 10% fetal bovine serum (FBS) at 37°C and 5% CO_2_. For drug treatment, HCT116 cells were treated with 5µM 5-Azacytidine for 48 hours, then washed with 1X PBS before harvesting.

### Crosslinking and Nuclei Preparation

Cells were grown to ∼75% confluency, harvested with trypsin, washed in 1× PBS, and frozen/stored at −80°C. Thawed cells were fixed in 1% formaldehyde and quenched in 0.125 M glycine, then washed twice in 1× PBS. Fixed cells were then resuspended in 500μl lysis buffer (50 mM Tris–HCl pH 8.0, 10 mM NaCl, 0.2% NP40, 1× PITC) for 30 min on ice with periodic resuspension. Lysed cells were spun 3500 RPM for 3 min and resuspended in 300 μl 1× NEB buffer 2, spun and resuspended in 198 μl 1× NEB buffer 2. 2μl of 10% SDS was added and incubated at 65°C for 10 min. After, 400 μl 1× NEB buffer 2 and 60 μl 10% Triton X-100 were added to quench the SDS. Samples were incubated at 37°C for 15 min. Nuclei were spun 3500 RPM for 3 min and resuspended in 300 μl 1× NEB buffer 2, repeat wash step.

### Protect-seq Protocol

Protect-seq protocol was performed as described in (Spracklin & Pradhan, 2020). Pelleted nuclei were resuspended in 183 μl DNaseI Buffer then 2 μl 100 mM Ca2+ (1 mM final), 5 μl DNase I (10 U), 5 μl MNase (10 000 U) and 5 μl RNase A (20 mg/ml) were added (200 μl final volume). Cells plus the enzyme cocktail were incubated at room temperature (RT) (also works at 37°C) for 30 min. Digested cells were spun 3500 RPM for 3 min and resuspended in 400 μl of 1× NEB buffer 2, then rotated at RT for 15 min. Digested/Wash#1 cells were spun 5000 RPM for 3 min and resuspended in the same 200 μl cocktail mix and incubated again at RT (or 37°C) for 30 min. Digested cells#2 were spun 10 000 RPM for 3 min and resuspended in 400 μl of 1× NEB buffer 2, then rotated at RT for 15 min (save aliquot for microscopy). Spin digested cells#2 10,000 RPM for 3 min and resuspended in 200 μl of 1× NEB buffer 2, 20 μl Proteinase K (SDS optional). Digest overnight at 65°C then purify using phenol/chloroform and ethanol precipitation (compatible with silica-bead purification).

### Illumina Library Preparation

DNA was quantified with Qubit (high-sensitivity) and sonicated using Covaris 50µl 300bp protocol. Illumina libraries were prepared using the NEB Ultra II DNA library kit using the manufacturer’s protocol. 4-5 PCR cycles were used to amplify NGS libraries and index samples.

### In situ Hi-C

HiC protocol was performed similar to (Rao et al., 2014). In brief, fixed nuclei were isolated, digested with MboI (NEB#R0147M), 5’ overhangs were filled-in with a biotinylated nucleotide, blunt-ends were ligated, followed by reverse crosslinking overnight. The purified DNA (2 µg) was sonicated using Covaris 50µl 400bp protocol. The sonicated DNA was brought to a volume of 400µl in binding buffer (5 mM Tris-HCl pH7.5; 0.5 mM EDTA; 1M NaCl) and mixed with 20µl of streptavidin magnetic beads (NEB#S1421) and rotated for 1hr at RT. The bead-bound DNA was washed twice with 400µl low-TE and resuspended in 50µl low-TE. NGS libraries were prepared using NEB DNA Ultra II kit (NEB#E7645). End prep: Mix 50µl of sample with 7µl End prep buffer and 3µl End prep enzyme, incubate for 30 min at RT then 30 min at 65°C, wash twice with 400µl low-TE and resuspended in 60µl low-TE. Adaptor ligation: 2.5µl of adaptor and 30µl of ligation mix was incubated at RT for 1-3hrs, washed twice with low-TE and resuspended in 90µl low-TE, following ligation, 3 µl USER was added for 30mins at 37°C, wash twice with 400µl and resuspend in 15µl. PCR: Add 5µl of universal F and index R primer, 25µl Q5 mix, 15µl sample for 5 PCR cycles. Libraries were purified with SPRI beads (0.9X) and quantified on a bioanalyzer and with NEB Illumina quant kit (NEB#E7630). HiC libraries were sequenced on a NextSeq500 either 150bp or 75bp paired-end reads.

### Chromatin Immunoprecipitation (ChIP) Experiments

SimpleChIP® Plus Enzymatic Chromatin IP Kit (Magnetic Beads) #9005 from Cell Signaling Technologies was used for all ChIP-seq experiments using the manufacturer’s recommended protocol. Four million cells per IP. Digested chromatin was pooled into a single tube for brief sonication to lyse nuclei. Supernatant was then split evenly between IPs (minus 2% input). Antibodies and chromatin were incubated overnight at 4°C, rotating. DNA was purified using spin columns and prepared using NEB Ultra II DNA library kit.

### Repli-seq

Repli-seq was performed and analyzed as described in (Marchal et al., 2018). In brief, cells were pulsed with 100uM BrdU for 2hrs, trypsinized, ethanol fixed, stained with propidium iodide and FACS sorted (SONY SH-800) based on DNA content (early S v late S). Genomic DNA was purified using Zymo DNA clean and concentrator and sonicated on a Covaris (S2) using the 300bp, 50ul protocol. Libraries were made with Ultra II DNA kits from NEB and sequenced on an Illumina miSeq and/or nextSeq.

## Computational Analysis

### Hi-C data processing

Hi-C libraries were trimmed with the *fastp* package (Chen et al., 2018) to remove low quality reads and sequencing adaptors. Hi-C datasets were processed using the *distiller* pipeline (https://github.com/open2c/distiller-nf) written for nextflow (Di Tommaso et al., 2017). Briefly, we mapped Hi-C sequencing reads to the human reference assembly hg38 using *bwa mem* (Li, 2013) with flags -SP. Alignments were parsed, filtered for duplicates and pairs were classified using the *pairtools* package (https://github.com/open2c/pairtools). Hi-C pairs were aggregated into contact matrices in the cooler format using the *cooler* package at multiple resolutions (Abdennur & Mirny, 2020). All contact matrices were normalized using the iterative correction procedure (Imakaev et al., 2012) after bin-level filtering.

### ChIP-seq and Protect-seq data processing

All ChIP-seq data, including data from (Lay et al., 2015) and (Maurano et al., 2015) but excluding those obtained from the ENCODE portal, were processed following the steps of the ENCODE ChIP-seq pipeline (https://github.com/ENCODE-DCC/chip-seq-pipeline2) with slight modifications using a simplified custom snakemake workflow. Briefly, reads were mapped to hg38 using *bwa mem* (Li, 2013). Alignment files (BAM format) were filtered for quality and duplicates using the *samtools* and *Picard* packages (Li et al., 2009). Cross-correlation analysis and fragment length estimation for single-ended datasets was performed using the *phantompeakqualtools* package (Landt et al., 2012). Signal track (target over input) generation was performed using *MACS2* (Y. Zhang et al., 2008). For CTCF, a motif instance was assigned to each ChIP-seq peak by scanning the core motif PWM (JASPAR MA0139.1) using *gimmemotifs* (Bruse & van Heeringen, 2018). Protect-seq data was mapped following the same procedure to produce signal tracks (treatment over input).

### Repli-seq data processing

Two-stage Repli-seq reads were processed following the protocol described in (Marchal et al., 2018). Replicates were merged to produce signal tracks of log_2_ count-normalized ratios of early divided by late fractions binned at 50-kb resolution. Tracks were then normalized by z-score transformation.

### Spectral analysis

To characterize long-range interaction profiles, 50-kb resolution Hi-C maps were dimensionally reduced by applying global eigendecomposition on *trans* contact frequencies. First, we manually identified and excluded three large translocated segments in HCT116 based on published karyotype analysis (Langer et al., 2005) narrowed down by visual inspection of Hi-C data in HiGlass (Kerpedjiev et al., 2018). Structural variations in DKO, on the other hand, were too widespread to systematically exclude so DKO clustering results were omitted from this study. Next, to mask the influence of *cis* data, we followed the same procedure described in (Imakaev et al., 2012), where *cis* pixels in the contact matrix are replaced with randomly sampled pixels from the same row or column. The resulting matrix was then re-balanced and scaled such that rows and columns sum to 1. Finally, the leading eigenvalues and associated eigenvectors of this matrix were then calculated using the eigsh routine from *numpy*, in descending order of eigenvalue modulus (i.e. not respecting algebraic sign). In order to standardize the spectrum of eigen*values*, eigendecomposition was done without a prior mean-centering step.

We describe our clustering method in more detail (see **Supplemental Note**). In summary, *m* leading eigenvectors were rescaled, concatenated as columns, and *k*-means clustering was applied to the rows using *scikit-learn*. We produced cluster assignments for a range of *k* for Hi-C maps of GM12878 (Rao et al., 2014), and unsynchronized untreated and 6h Auxin-treated Rad21-AID HCT116 (Rao et al., 2017), calculated silhouette scores (**S.Figure 1**) and visually compared cluster profiles to a large number of independent genomic tracks. The final number of clusters was chosen based on a balance of clustering metrics and interpretability.

For visualization of the approximate manifold structure, further dimensionality reduction on the *m* leading eigenvectors was performed using UMAP (McInnes et al., 2018). Additionally, direct visual inspection of the unreduced eigenvector subspaces (pairwise) and related genomic and functional data proved to be indispensable for interpretability of clusters (see section on shaded scatter plots).

### Raster (shaded) scatter plots

The new *matplotlib* (Hunter, 2007) extension for the data graphics pipeline *datashader* (dsshow function) (https://datashader.org) was used to generate scatter plot visualizations of points representing 50kb genomic bins. The datashader pipeline is used to prevent overplotting a dense point cloud by aggregating points onto a regular 2D grid and either (i) color-mapping the resulting raster to associated quantitative values (e.g. point count, mean value) or (ii) displaying associated color-coded categorical values (cluster labels, chromosome, etc.) via image compositing.

### ChromHMM state assignment

We ran *ChromHMM* (Ernst & Kellis, 2012) to create epigenomic segmentations for HCT116 and DKO using bam files for ChIP-seq of broad marks/factors HP1a, HP1b, H3K9me3 and H3K27me3. For HCT116, we also included data for SON TSA-seq (Zhang et al., 2020). Tracks were binarized at 50kb using BinarizeBam and were modified to ignore bins filtered in Hi-C data. Models were trained using 50kb bins (LearnModel -b 50000) for a range of state numbers. A 7-state model was chosen for HCT116. For DKO, a 6-state model was able to qualitatively capture the same repressive states based on emission parameters (with only a single active state, since TSA-seq was not available to discriminate between two active states).

### Chromatin state analysis

Gene quantification table for HCT116 was obtained from ENCODE and cross-referenced to GENCODE v29 basic gene annotations for hg38. Records were intersected against interaction profile group labels using *bioframe* and grouped. Adjusted TPM values were log-transformed and binned to generate histograms. Kernel density estimates were generated using *seaborn* (Waskom, 2021).

HCT116 and DKO Whole Genome Bisulfite sequencing data (hg19) from (Blattler et al., 2014) were lifted over to hg38 using *Crossmap* (Zhao et al., 2014). DNA methylation tracks for HCT116 and 5Aza-treated cells (24h) generated using Hybrid Selection Bisulfite Sequencing (hg19) from (Johnstone et al., 2020) were also lifted over to hg38 using *Crossmap*. All data were filtered for CpG context to exclude liftover base changes. A custom script was used to aggregate records into 50-kb bins and calculate the cumulative methylation fraction from CpGs divided by total number of CpGs per bin.

Functional profiles for spectral clusters (as in Fig 1D, and averages in 2B) were derived from categorical or mean-aggregated quantitative signal tracks (distance from centromere, LaminB1 DamID-seq, SON TSA-seq, Protect-seq, Repli-seq, WGBS, ChIP-seq) at 50-kb resolution to match the resolution of interaction profile analysis.

Interaction profile domain metaplots and stacked signal heatmaps were generated from BigWig files using the *pybbi* package (https://github.com/nvictus/pybbi). Unscaled stacked heatmaps were defined using the domain midpoints as a reference point flanked by a fixed genomic distance left and right, while re-scaled stacked heatmaps were generated by independently partitioning the intradomain signal and flanking regions into a fixed number of bins. Metaplots were generated by averaging re-scaled heatmaps vertically.

Sankey plots were generated by using ChromHMM segmentation maps from DKO cells. Chromatin states were intersected against disrupted domains using *bioframe*. Next, total base pairs overlapped for each chromatin state were counted. Sankey plots were generated using *plotly*.

### ROC curves

To assess the correspondence of individual signal tracks to interaction profile group assignments derived from Hi-C data, we treated each mean-aggregated 50-kb resolution track as a binary classifier to predict a given interaction profile label (one of A_1_, A_2_, B_0_, B_1_, B_2_) by applying a simple value-based discrimination threshold on the signal track. ROC curves and area under ROC for these classifiers were calculated using *scikit-learn*. Curves that dip below the diagonal indicate thresholds with predictive power for the complement of the target label (e.g. not A_1_).

### Quantile-based ChIP-seq and Hi-C histograms

50-kb resolution ChIP-seq tracks were grouped into percentiles of either E1 signal or H3K9me3 signal to generate histograms and standard deviation envelopes.

Expected contact frequency profiles were generated using *cooltools* (https://github.com/open2c/cooltools) and bivariate histograms of observed/expected contact frequency (a.k.a. saddle plots) using percentiles of either E1 or H3K9me3 signal as bins were also generated using *cooltools*.

### H3K9me3 domain calling

Domains defined by broad H3K9me3 ChIP-seq enrichment across six cell types (HCT116, HFFc6, IMR90, K562, GM12878, H1-hESC) were called using a hidden Markov Model procedure. H3K9me3 ChIP-seq bigwigs were mean-aggregated at 25-kb, log-transformed and z-scored, and binarized with a threshold of 1, which were used to train a 2-state Bernoulli HMM using *pomegranate*. Smoothed runs of 1s from the Viterbi parses were used to define domains.

### P(*s*) curves per interaction profile group

Scaling curves of contact frequency *P* as a function of genomic separation *s* were generated using *cooltools* by aggregating normalized contact frequency over valid pixels along diagonals of 1kb-resolution *cis* contact maps limited to interaction profile domains, with diagonals grouped into geometrically increasing strata of genomic separation. Average contact frequency *P(s)* curves are displayed using log-log axes.

### Insulation analysis

Diamond insulation scores (Crane et al., 2015) were calculated on 25kb resolution Hi-C maps with a 100-kb sliding window using the *cooltools* package. Additionally, an insulation minimum calling procedure based on peak prominence, described in (Nora et al., 2017) was used to call insulating loci from the insulation score signal.

### Hi-C pileup maps

The *cooltools* package was used to calculate aggregate observed-over-expected contact frequency maps (pileup maps) centered at CTCF sites and bounded by a fixed flanking genomic distance. Pileup maps are centered on the main diagonal at each feature’s midpoint.

### Replication timing domain analysis

To identify early and late replicating domains, a 25-kb binned pandas dataframe was generated using *bioframe*. HCT116 and DKO replication timing signal tracks were imported into the binned dataframe using *pybbi*. Missing values were represented as NaN. Domains were identified with a 2-state Gaussian HMM using Pomegranate (Schreiber, 2017). Viterbi state calls were made on a per bin basis and used for downstream analysis. Neighboring states were merged to create domains then converted to bed files (https://github.com/gspracklin/hmm_bigwigs).

Differential replication timing loci were identified by applying a cutoff of 0.75 on the difference between HCT116 and DKO 50-kb z-score tracks. Differentially timed loci separated by up to 250 kb were then merged into larger intervals using bioframe.cluster to produce 199 differentially timed regions.

### Polymer simulations

Simulations were created using the Polychrom library (Imakaev et al., 2019). The polymer simulations ran using the OpenMM engine for GPU-assisted molecular dynamics simulations (Eastman et al., 2017). Each simulation modeled 8-11 megabases of chromatin fiber as a chain of 1-kilobase monomers, and included five copies of the system inside the same container. Each simulation was run for 500,000,000 molecular dynamics steps. Periodic boundary conditions were used to maintain a density of 0.2 monomers per cubic nanometer.

The following energies are in terms of kT, and distances are measured in terms of the diameter of the monomers, which is 20 nanometers. Adjacent monomers on the chain are connected by a harmonic bond with potential U = 100(r−1)^2^. Polymer stiffness is modeled by U = S(1−cos(α)), a force dependent on the angle α formed by three adjacent monomers, and S is a stiffness parameter equal to 1.5.

To model loop extrusion, LEFs were probabilistically loaded onto the polymer chain at uniformly random positions. Each LEF is represented by a harmonic bond equivalent to the one that connects adjacent monomers on the chain. Each step of 1D dynamics corresponded to 400 molecular dynamics steps. An LEF with an upstream leg at monomer *i* will stay at *i* with probability ½ and move to *i*-1 with probability ½ each step, unless *i*-1 is occupied by an LEF or a CTCF. Similarly, a downstream leg at monomer *j* will stay at *j* with probability ½ and move to j+1 with probability ½, unless *j*+1 is occupied by an LEF or CTCF. CTCF sites were placed at fold-change peaks in HCT116 CTCF ChIP-seq (ENCODE ID ENCFF549PGC), with directionality according to CTCF motifs (from (Maurano et al., 2015)). Each CTCF had a capture probability of min((fc-1)/fc_med,1), where fc is the CTCF fold change and fc_med is the median CTCF fold change over the region. Legs were released from CTCFs with a probability of 0.006 each monomer step. Each LEF was unloaded with a probability of 1/100 each step of 1D dynamics, and LEFs were separated by an average of 600 monomers.

