## Supplemental Note for "Heterochromatin diversity modulates genome compartmentalization and loop extrusion barriers"

### Spectral clustering of Hi-C data

Here we describe a spectral clustering algorithm to identify loci with common long-range contact frequency profiles in Hi-C data. As in previous work, we focus on interchromosomal (*trans*) contact frequencies. Because *trans* contacts are experimentally sampled less frequently than intrachromosomal (*cis*) contacts, this strategy works best on deeply sequenced contact maps with strong pattern contrast in *trans*. However, it provides the advantages of not having to detrend intra-arm and inter-arm polymer scaling relationships and working on more than one chromosomal arm at a time. For example, [1] used contact frequencies between two sets of disjoint chromosomes (odd and even-numbered) and clustered *trans* Hi-C data along each axis (rows and columns) separately at 100-kb resolution, followed by harmonizing cluster identities between the two axes.

Clustering leading eigenvectors or principal components (i.e., spectral clustering) provides a scalable and noise-reducing alternative to clustering complete contact matrices directly. In [2], this was done separately on each chromosome using *cis* data, followed by a recursive clustering step using *trans* data to harmonize cluster identities between chromosomes. In this study we apply dimensionality reduction using global eigendecomposition on *trans* contact frequencies from genome-wide balanced 50kb-resolution Hi-C maps. This provides a single collection of eigenvectors for clustering, requiring no harmonization between chromosome arms or sets thereof. As an additional benefit, it also facilitates further dimensionality reduction (embedding) and visualization.

Before eigendecomposition, because *trans* translocations contaminate interaction profiles and will exert an influence on higher order eigenvectors, we excluded such regions from analysis. We manually identified and excluded three large translocated segments in HCT116 (chr8: 67.35 Mb – end; chr16: 78.93 Mb – end; chr17: 43.40 Mb – end) based on published karyotype analysis [3] narrowed down by visual inspection of Hi-C data in HiGlass [4]. Unfortunately, structural variations in DKO were too widespread to systematically exclude, so DKO clustering results were omitted from this study. To mask the influence of *cis* data, we followed the same procedure described in [5], where *cis* pixels in the contact matrix are replaced with randomly sampled *trans* pixels from the same row or column.

The resulting matrix was then re-balanced and scaled such that rows and columns sum to 1. In order to standardize the spectrum of eigenvalues, eigendecomposition was done without the prior mean-centering step used in [5]. The decomposition of this $n \times n$ normalized affinity matrix $A$ (interpreted as a weighted graph) can be expressed as a sum of rank-1 outer products:

$$A = \lambda_0 \vec{e}_0 \vec{e}_0^T + \lambda_1 \vec{e}_1 \vec{e}_1^T + \lambda_2 \vec{e}_2 \vec{e}_2^T + \ldots + \lambda_{n-1} \vec{e}_{n-1} \vec{e}_{n-1}^T, \quad (1)$$

where $\lambda_0 = 1$, $\vec{e_0} = \frac{1}{\sqrt{n}} \mathbb{1}$, $||\vec{e_i}|| = 1$ and

$$1 > |\lambda_1| > |\lambda_2| > \ldots > |\lambda_{n-1}| > 0$$

The largest eigenvalue $\lambda_0$ is 1 and its multiplicity depends on the number of connected components of the graph whose edge weights are given by $A$. Because a Hi-C matrix of sufficient coverage represents a single connected component (ignoring filtered bins), $\lambda_0$ is uniquely equal to 1 and is associated with a constant-valued eigenvector $\vec{e}_0$ that may be discarded for clustering purposes [6, 7]. The remaining eigenvalues lie in the range $(-1, 1)$ and those with relatively large modulus are most informative for clustering [8, 9]. Interestingly, the next leading eigenvector ($\vec{e}_1$) is identical to the usual A/B compartment eigenvector and also happens to be the well-known "Fiedler vector" in spectral clustering [7]. Hence, the classical bipartitioning of genomic loci based on $\vec{e}_1$ (or equivalently, the first principal component using PCA) happens to be a special case of spectral clustering.

To extend this approach in order to obtain a finer and more informative partition, we selected the $m$ leading eigenvectors after $\vec{e}_0$ ($m < n-1$). The leading eigenvalues and associated eigenvectors of $A$ were calculated using the `eigsh` routine from *numpy* [10], in descending order of eigenvalue modulus (i.e. not respecting algebraic sign). The approximation can be written as

$$\begin{aligned} A &\approx \vec{e}_0 \vec{e}_0^T + \lambda_1 \vec{e}_1 \vec{e}_1^T + \ldots + \lambda_m \vec{e}_m \vec{e}_m^T \\ &= \vec{e}_0 \vec{e}_0^T + \\ &\quad \mathrm{sign}(\lambda_1) \sqrt{|\lambda_1|} \vec{e}_1 \cdot (\sqrt{|\lambda_1|} \vec{e}_1)^T + \ldots + \\ &\quad \mathrm{sign}(\lambda_m) \sqrt{|\lambda_m|} \vec{e}_m \cdot (\sqrt{|\lambda_m|} \vec{e}_m)^T \end{aligned} \quad (2)$$

Motivated by equation (2), the individual unit-normed eigenvectors (excluding $\vec{e}_0$) were weighted as shown ($\sqrt{|\lambda_1|}\vec{e}_1$, $\sqrt{|\lambda_2|}\vec{e}_2$, ..., $\sqrt{|\lambda_m|}\vec{e}_m$), concatenated as columns, and $k$-means clustering was applied to the rows using *scikit-learn*'s `KMeans` estimator [11]. The estimator was run with 100 centroid initializations using the `kmeans++` initializer, which returned the best run in terms of inertia. As a heuristic, $m$ was chosen to be the maximum number of eigenvalues before the first negative eigenvalue appeared (i.e., before succeeding eigenvalues began oscillating around 0), though empirically for a given $k$, clustering was observed to be stable beyond that number as long as genomic regions undergoing translocations were excluded from analysis. We produced cluster assignments for a range of $k$ for GM12878 [1] and both unsynchronized untreated and 6h Auxin-treated Rad21-AID HCT116 maps [12], calculated silhouette scores and visually compared cluster profiles to a large number of independent genomic tracks. The final number of clusters for HCT116 ($k = 8$) was chosen based on a balance of clustering metrics and interpretability.

Finally, we note that readers familiar with spectral partitioning of graphs may be more accustomed to discussions of the eigenvectors of the graph Laplacian matrix rather than the affinity matrix $A$. The Laplacian is $L = D - A$, where $D$ is a diagonal matrix with the degrees $d_1, \ldots, d_n$ on the diagonal, and its most common normalized forms used for spectral clustering are $L_{sym} = D^{-1/2}LD^{-1/2}$ and $L_{rw} = D^{-1}L$ [7]. Note that in our case, $A$ is a stochastic matrix (i.e. a graph with constant degree 1) and its (normalized) Laplacian is simply $I - A$, which has the exact same eigenvectors as $A$ and shifted eigenvalues $\lambda'_i = 1 - \lambda_i$.
